# Turn ‘noise’ to signal: accurately rectify millions of erroneous short reads through graph learning on edit distances

**DOI:** 10.1101/2024.04.05.588226

**Authors:** Pengyao Ping, Shuquan Su, Xinhui Cai, Tian Lan, Xuan Zhang, Hui Peng, Yi Pan, Wei Liu, Jinyan Li

## Abstract

Although the per-base erring rate of NGS is very low at 0.1% to 0.5%, the percentage/probability of *erroneous reads* in a short-read sequencing dataset can be as high as 10% to 15% or in the number of millions. Correction of these wrongly sequenced reads to retrieve their huge missing value will improve many downstream applications. As current methods correct only some of the errors at the cost of introducing many new errors, we solve this problem by turning erroneous reads into their original states, without bringing up any non-existing reads to keep the data integrity. The novelty of our method is originated in a computable rule translated from PCR erring mechanism that: a rare read is erroneous if it has a neighbouring read of high abundance. With this principle, we construct a graph to link every pair of reads of tiny edit distances to detect a solid part of erroneous reads; then we consider them as training data to learn the erring mechanisms to identify possibly remaining hard-case errors between pairs of high-abundance reads. Compared with state-of-the-art methods on tens of datasets of UMI-based ground truth, our method has made a remarkably better performance under 19 metrics including two entropy metrics that measure noise levels in a dataset. Case studies found that our method can make substantial impact on genome abundance quantification, isoform identification, SNP profiling, and genome editing efficiency estimation. For example, the abundance level of the reference genome of SARS-CoV-2 can be increased by 12% and that of Monkeypox can be boosted by 52.12% after error correction. Moreover, the number of distinct isomiRs is decreased by 31.56%, unveiling there are so many previously identified isomiRs that are actually sequencing errors.

**Author summary:** Detecting short-read sequencing errors and correcting the related erroneous reads is a long-standing problem in bioinformatics. Current error correction algorithms correct only small parts of the errors but simultaneously introduce thousands of non-existing sequences. We present a new method to rectify erroneous reads under 300 bp produced by PCR-involved miRNA-sequencing, small RNA sequencing, or paired-end RNA sequencing, regardless of platform or sample type. Our method is the first kind considering the PCR erring mechanism and machine learning technique to improve sequencing data quality by turning millions of erroneous short reads into their original state without bringing up any non-existing sequences into the read set. Our error correction method can make a significant impact on a wide range of cutting-edge downstream applications. The observations and advantages in the case studies lay down strong evidence to question the accuracies of current downstream research outcomes and open new avenues to conduct downstream analysis whenever short-read data are adopted.

## Introduction

Next-generation sequencing (NGS) techniques and platforms have dramatically changed the world of genomics and computational biology [1–3]. High throughput DNA-sequencing has enabled large scale whole-genome sequencing and gene-targeted sequencing; NGS-based RNA-seq has provided ever higher coverage and sharper resolution of dynamic transcriptome for a wide range of applications such as isoform discovery, differential gene expression analysis, alternative gene splicing and allele-specific expression profiling [2]. However, NGS inevitably self-made sequencing errors including base deletions, insertions and substitutions at various steps like sample handling, library preparation, polymerase chain reaction (PCR), and/or at the base calling step [4, 5]. Although the erring rate is estimated very low at 0.1% - 0.5% per base in Illumina short-read sequencing, huge numbers of erroneous bases have been generated and stored at every sequencing dataset (e.g., about 197,402 base errors in a miRNA-sequencing dataset ERR187525, and about 997,020 base errors in a paired-end whole-genome sequencing dataset SRR22085311 which have been found through this study). As these mistaken bases are randomly distributed across all the reads in a dataset, the percentage/probability of *erroneous reads* in a dataset can be very high (e.g., as high as 10-15%).

Suppose the per-base erring probability is estimated as *p* at a sequencing platform, and assume these erring events are independent at all the base positions in a read, then the probability *p*_*error*_(*r*) of a read *r* containing one or multiple base errors is given by

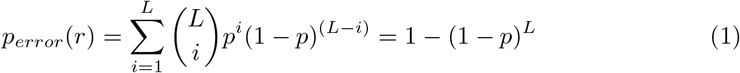

where, *L* = ∥*r*∥, the length of read *r*. If *p* = 0.1% and *L* = 100, then *p*_*error*_(*r*) = 9.52%. In other words, the percentage of erroneous reads in a dataset is about 9.52% when the per-base erring rate is estimated as 0.1% and the length of reads *L* = 100bp. If the per-base erring rate *p* is estimated as 0.15%, then there are about 13.94% of erroneous reads in the dataset.

This is a fundamental issue previously unrecognized concerning the high percentages of erroneous reads in NGS datasets. These erroneous reads are usually treated as data noise implicitly or explicitly excluded for downstream data analysis such as de novo genome/transcriptome assembly and differential gene expression profiling [6, 7]. Or, these erroneous reads are un-purposely considered as genuine true reads in the data analysis which may have led to inaccurate or wrong conclusions. To restore the huge missing value of these high percentages of erroneous reads in each sequencing dataset, it is highly demanded to do accurate rectification of all these errors, as opposed to treating them as noise removal, to boost the data quality and integrity so as to improve the downstream applications.

One of the main sources of the sequencing errors is from PCR (polymerase chain reaction), a technique that makes fast duplications of small segments of DNA which has been used by NGS to amplify the fragmented DNA/RNA molecules for effective sequencing. Most of the time, PCR makes perfect copies of the fragmented segments of DNA/RNA, but occasionally it introduces base-pair substitutions, deletions, insertions, or even yields new hybrid sequences during template switching [8]. Thus, after the PCR amplification, one or two copies in the duplications of a DNA segment may show inconsistent bases. Fig. 1a illustrates how base errors arise when amplifying one DNA template during PCR amplification. PCR errors not only occur in the library preparation but also during sequencing processes such as clonal molecules [5]; Fig. 1b is an example that depicts how errors are introduced in the process of bridge amplification during Illumina sequencing. Such PCR erring incidents are then inherited by NGS’s base calling step that converts the nucleotide sequence into a digital string named a read. The conversion is not 100% accurate as well, similar to PCR introducing minor mistakes (Fig. 1c) [4, 9]. So, there exist various erring incidents in the sequencing.

**Fig 1.**
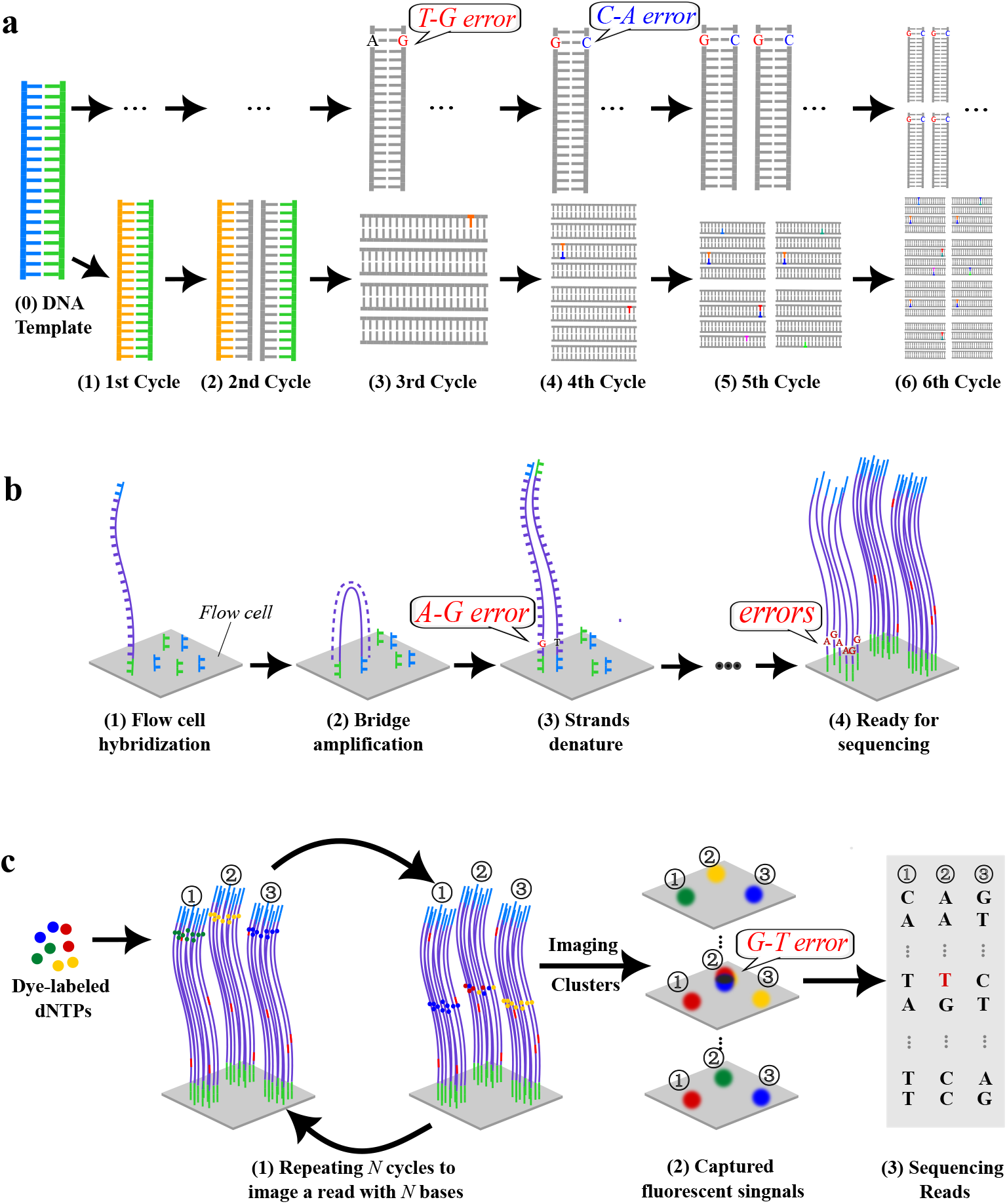
Schematic diagrams to illustrate how base errors are generated during library preparation and sequencing process. (a) Schematic illustration of base error generation when amplifying one DNA template during conventional PCR amplification. Base ‘T’ mutated to ‘G’ in the 3rd cycle, the ‘C’-to-’A’ error occurs in the 4th cycle and this error is inherited by the subsequent cycles. (b) Schematic graph depicting PCR errors generated in the process of bridge amplification during Illumina sequencing. An example of ‘A’-to-’G’ is inherited. (c) An overview of base calling during Illumina sequencing.

However, one thing nearly certain is that: an erroneous read must be a low-frequency rare read if the error occurred at the late cycles of PCR. This is because the probability of the same mistakes happening at the same positions is extremely low, especially in 200-300bp reads.

Efficient detection of these erroneous reads from a dataset of hundreds of millions of reads is challenging. First, some low-frequency rare reads are genuine reads not containing any sequencing errors. This is attributed to the uneven PCR amplification rates at different segments of the DNA—poorly amplified molecules will be sequenced to a lesser extent than the highly amplified molecules and vice versa [10, 11]. Second, an amplified segment after PCR erring may become identical to a high-frequency molecule. As a result, for a pair of high-frequency reads A and B that are very similar, it is impossible to judge, without machine learning of the PCR erring mechanisms, whether A’s copies contain B’s wrongly amplified copy, or whether B’s copies contain A’s wrongly amplified copy.

We construct a graph *rG*(*R*) using the unique reads *r*_1_, *r*_2_, …, *r*_*n*_ along with their respective frequencies from a reads dataset *R* (a multiset of reads) to detect erroneous reads under the sophisticated help of graph-based machine learning. Let *freq*(*r*) represent the abundance level or the frequency of a read *r*, or the number of copies of *r* in the sequencing data. For each of the unique reads in *R*, we represent it as a node in the graph and label the node with the read’s frequency. There is an edge *e*_(*i,j*)_ between node *r*_*i*_ and node *r*_*j*_ if the *edit distance* between read *r*_*i*_ and read *r*_*j*_ is 1 or 2.

Specifically, when searching for edges with an edit distance of 2, only substitutions are taken into account. A read *u* is a neighbouring read of read *v* if there is an edge between them. As understood from the PCR erring mechanism in NGS, the pairing of two neighbouring reads *u* and *v* implies that a copy of *u* is a wrongly amplified/sequenced copy of the *v* molecule, or a copy of *v* is a wrongly amplified/sequenced copy of the *u* molecule, or both. When *freq*(*v*) is low while *freq*(*u*) is high, we rectify the erroneous read *v* by removing this node from the graph, while increase *freq*(*u*) by *freq*(*v*). That is, we turn the ‘noise’ read *v* (low-frequency rare read) into its normal state *u*. We denote such a set of erroneous reads in the graph as `edit-erring-READS` and the isolated nodes with high frequencies as `error-free-READS`.

Using a small edit distance of 1 or 2 to define the edges of the graph is because those erroneous reads containing one mistaken base or two constitute the majority of the total erroneous reads in the dataset. The majority percentage is given by

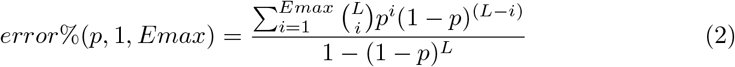

where *p* is the base erring probability, *Emax* is a maximum edit distance allowable to define an edge. If *L* = 100, *p* = 0.1% and *Emax* = 2, then *error*%(*p*, 1, *Emax*) = 99.84%. This indicates that 99.84% of all the possible erroneous reads in the dataset are those reads containing one base error or two (Emax).

The second challenge in the correction of erroneous reads in the graph *rG*(*R*) is to deal with the situation when a low-frequency read is linked to multiple high-frequency reads, or two (or more) high-frequency reads are linked each other in the graph (denoted as ambiguous errors). Hence, we model the situations as a classification problem and use machine learning techniques to predict whether the duplications of a high-frequency read contain or not contain wrongly sequenced copies of its neighbouring high-frequency reads.

This is a novel classification problem not formulated in any literature. In this work, we use `edit-erring-READS` and `error-free-READS` as training data and extract multiple features of different dimensions from the data, and then utilize an optimized gradient boosting classifier of the eXtreme Gradient Boosting (XGBoost) [12] to make the prediction under a supervised learning framework. As the training data is *rG*(*R*)-specific, the prediction model is able to learn the inherent erring patterns of each specific sequencing platform that conducts the specific biomolecular samples’ sequencing. So, our machine learning approach is competent to handle the rectification of erroneous reads that have a length less than 300 bp produced by any PCR-involved miRNA-sequencing, small RNA-sequencing, or paired-end RNA-sequencing regardless of the difference in the platforms or in the biomolecular samples. We name our method ‘noise2read’.

## Results

Comparing with state-of-the-art methods including *k*-mer-methods [13–19], multiple sequence alignment based methods [20–24], and other methods [25–27], our noise2read consistently outperforms under 19 metrics on eight UMI-based wet-lab datasets and five simulated single-end and paired-end datasets constructed in this study. It also has superior performance on eight UMI-based wet-lab datasets and four simulated miRNA datasets established previously in published literature. Case studies on twelve single-end and two paired-end sequencing datasets (about abundance change of viral reference genomes, isoform identification, SNP profiling, genome base editing efficiency estimation) reveal that noise2read can make substantial impacts on downstream applications. We first present real distributions of the erroneous reads containing various numbers of base errors in a dataset through UMI-based cluster analysis.

### High prevalence of erroneous reads containing one or two base errors from UMI-based cluster and distribution analysis

We utilized the sequence information of UMI tags to investigate the distributions of erroneous reads that contain different numbers of base editing errors. Specifically, we divided the reads in a UMI group into high-frequency reads and low-frequency reads. Then, we calculated the edit distance between each unique low-frequency read and each unique high-frequency read. Then, each of the unique low-frequency read has the smallest edit distance with the set of unique high-frequency reads. Given each of these smallest distances, we record the number of low-frequency sequences that have this edit distance with the set of high-frequency reads.

We applied the above process to the datasets of SRR1543964-SRR1543971 by defining a high-frequency read as a read with a copy count no less than five (note: clusters with ambiguous base ‘N’ in the UMI sequence are not included in this analysis). We observed that there exist two different high-frequency sequences that have been tagged with the same UMI, as similarly reported in the literature [28]. For instance, as seen in S1 Fig, each of these UMI clusters has two high-frequency reads, between which the edit distance is bigger than 100 (111 or 129 respectively), demonstrating that such two high-frequency reads within the same UMI cluster should be originated from two different molecules although they were tagged with the same UMI.

Moreover, there exist big editing distances (e.g., 116) between high and low-frequency reads within the same UMI cluster, it would be unreasonable to assume only base-editing-error-relationship between all the low and high frequency reads. In fact, a low-frequency sequence with a small edit distance to a high-frequency read is more likely caused by PCR or sequencing errors. Here, we assume that those low-frequency reads with an edit distance equal to or less than 4 may be erroneous reads caused by PCR or sequencing errors. In this context, among these eight data sets, at least 60% of the erroneous reads in 95.21%-96.70% of the UMI clusters are caused by 1 or 2 base errors, as depicted in the stacked bar chart in S2 Fig. Five more UMI clusters are shown in S3 Fig: 81.25%, 95.24%, 100%, 94.73% and 100% of low-frequency reads have the 1 and 2 base difference with the set of high-frequency reads in the same UMI cluster. These findings indicate that those erroneous reads containing one mistaken base or two constitute a more significant proportion of the total erroneous reads in the data set. Based on our theoretical analysis and UMI cluster analysis, the proposed algorithm, noise2read, is set to correct erroneous reads containing base errors less than 3.

### Constructed training data for noise2read

Noise2read is a progressive three-stage error correction method. As introduced above, its novelty sits in the computable rule translated from PCR erring mechanism: a rare read is erroneous if it has a neighboring read of high abundance. With this principle, we construct a graph to link every pair of reads of a small edit distance to detect a substantial part of erroneous reads in the graph. Then we take them as training data to learn the platform-specific erring mechanism to identify possibly remaining hard-case errors between pairs of frequent reads in the graph, namely specific training data is used at different platforms.

An Auto Machine Learning (AutoML) module is centered in the process of noise2read, which is used multiple times in the different stages for the prediction of ambiguous or amplicon errors. AutoML has a component for the preparation of training and objective data and has a component for the parameter optimization of the gradient boosting-based classifiers (Fig. 2). The first stage (shaded in blue) rectifies low-frequency leaf nodes (genuine errors) and ambiguous errors by a traversal on the 1nt-edit-distance read graph 1*nt*-*rG*(*R*_0_) constructed from the original reads dataset *R*_0_. Here, every edge in the 1nt-edit-distance read graph means the edit distance between the two nodes is one nucleotide (i.e., 1nt). The second stage (shaded in pink) conducts correction of genuine and ambiguous errors at the 2nt-edit-distance read graph 2*nt*-*rG*(*R*_1_) constructed from the first-stage corrected dataset *R*_1_. Here, every edge in the 2nt-edit-distance read graph means the edit distance between the two nodes is two nucleotides (i.e., 2nt). Particularly, we consider only substitution relationships for constructing 2nt-edit-distance edges since the majority of NGS data conforms to a consistent read length. The third stage (shaded in yellow) is designed to eliminate specific errors at an updated 1nt-edit-distance graph 1*nt*-*rG*(*R*_2_) only for the amplicon sequencing data but using the same AutoML module for prediction.

**Fig 2.**
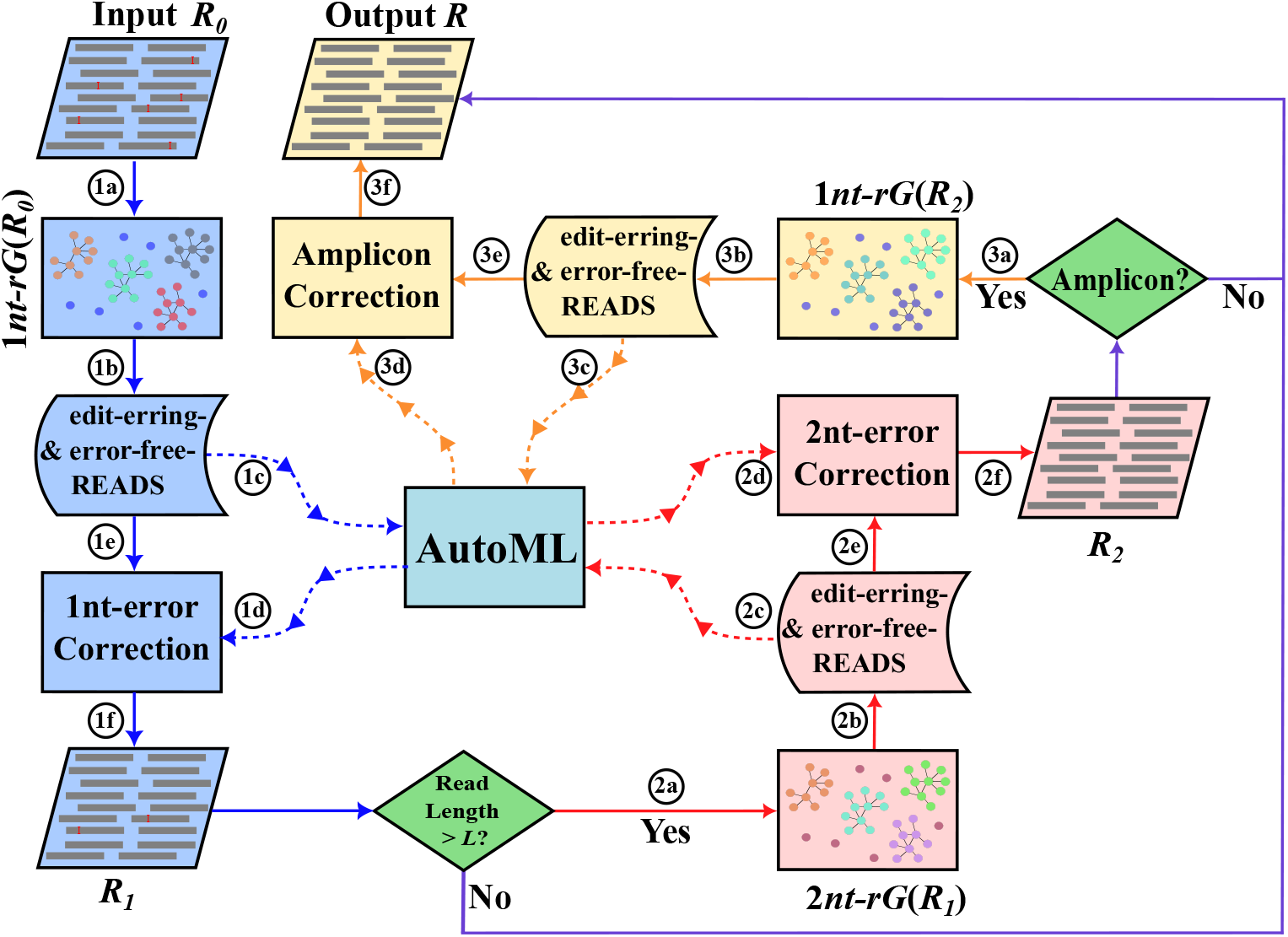
Overview of the workflow of noise2read. The first stage 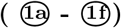 and the second stage 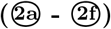 rectify 1nt- and 2nt-based-errors to their normal states, respectively. The third stage 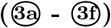 is optional only for further correction specified to the amplicon sequencing data. The integrative Auto Machine Learning (AutoML) module is used multiple times by feeding different `edit-erring-READS` and `error-free-READS` in each stage.

Graph *rG*(*R*) is often a disconnected graph. Fig. 3 shows nine sub-graphs of *rG*(*D*1) constructed at the first stage, where *D*1 is a simplified version of SRR1543964. It is interesting to see that there are many clustered low-frequency leaf reads linked to one high-frequency read, while there also exist edges that link pairs of high-frequency reads.

**Fig 3.**
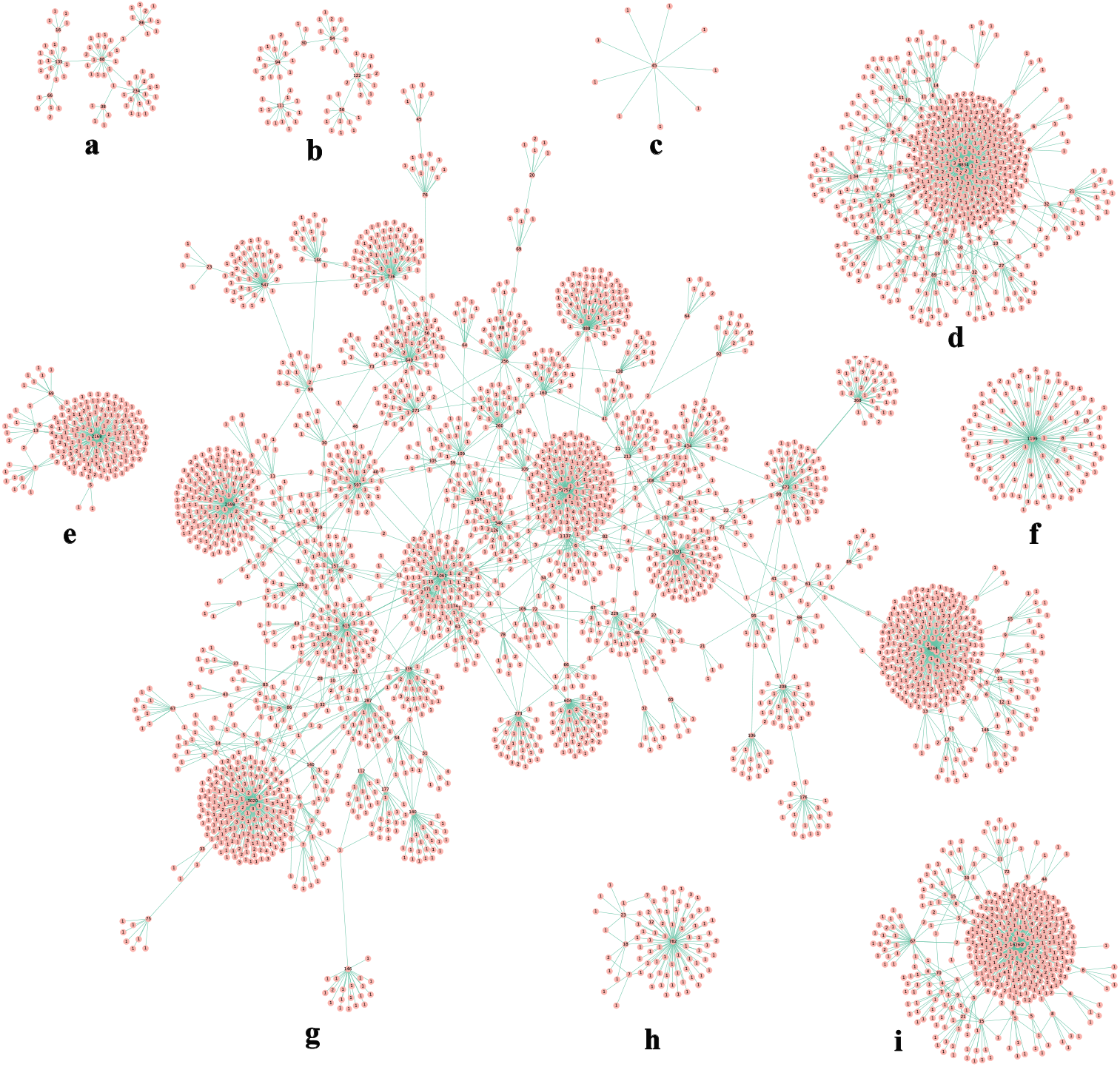
Nine subgraphs, denoted as (a-i), from 1*nt*-*rG*(*R*_0_) in the first correction stage for the dataset *D*1 derived from SRR1543964. Each node in the subgraphs represents a read and is labelled with its frequency inside the circle. The edge connecting two nodes indicates that their edit distance equals 1. The number of nodes in the subgraphs (a, b, c, d, e, f, g, h) or (i) is 79, 69, 10, 621, 190, 111, 2788, 83 or 445, respectively.

Fig. 4 is a zoomed version with more details about a subgraph in Fig. 3a, where the high-frequency nodes are highlighted in orange and the low-frequency nodes are highlighted in pink. By definition, every edge in this graph implies that the linked reads have only one base difference. With these sub-graphs, noise2read

**Fig 4.**
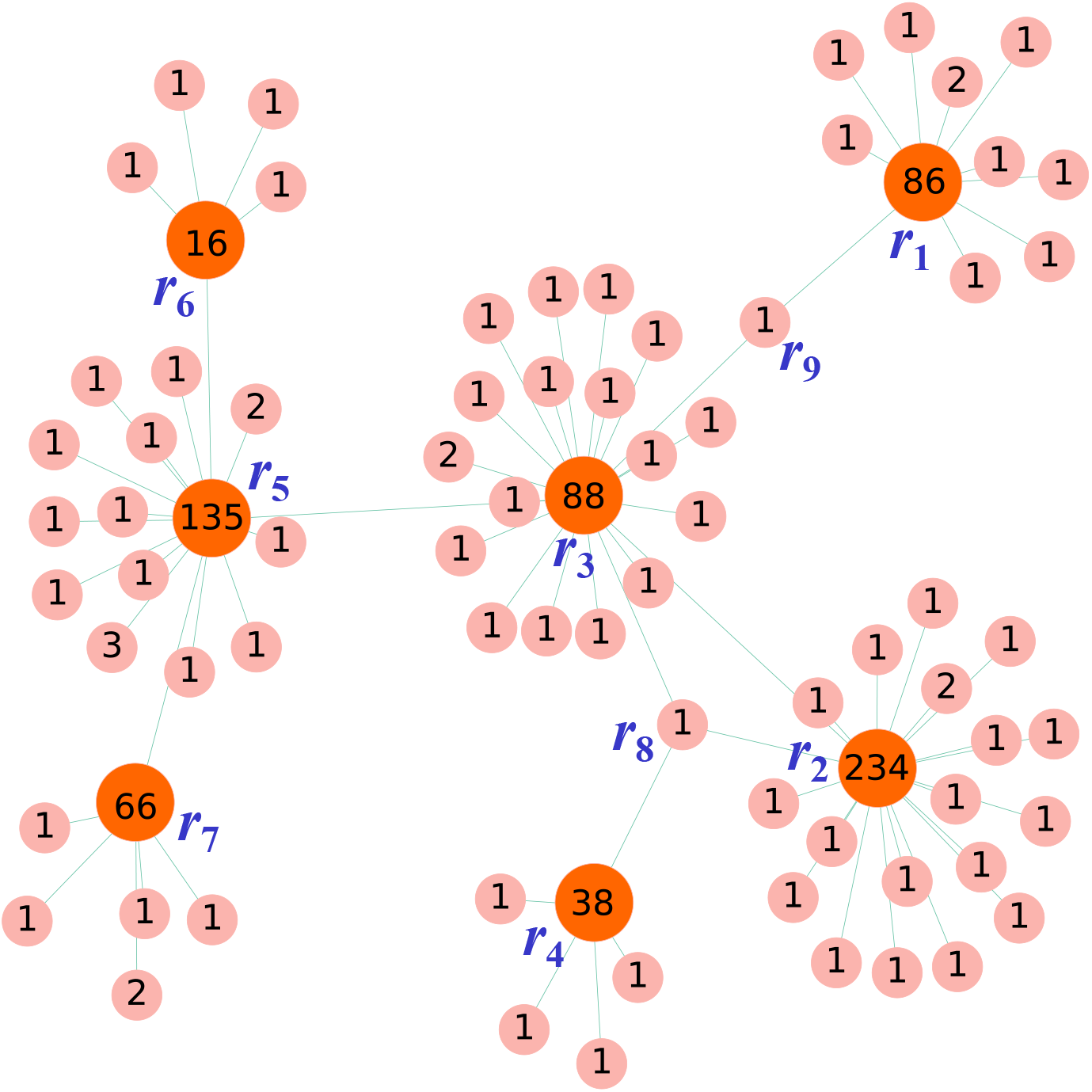
This subgraph is a zoomed-in version of Figure 3a; it contains six high-frequency reads labelled as *r*_1_ to *r*_7_ and highlighted in orange, as well as contains 72 low-frequency reads shaded in pink.

- directly turns those leaf nodes of low-frequency into their high-frequency parent nodes (their normal states *r*_1_, *r*_2_, …, or *r*_7_);
- uses the AutoML module to identify the parent node of two low-frequency nodes *r*_8_ and *r*_9_, as these two low-frequency nodes are each linked to more than one high-frequency read (*r*_8_ is linked to *r*_2_, *r*_3_ and *r*_4_; *r*_9_ is linked to both of *r*_1_ and *r*_3_); and
- uses the AutoML module to judge whether there are erroneous reads between the linked high-frequency nodes (e.g., between *r*_2_ and *r*_3_, between *r*_5_ and *r*_7_).

Although noise2read is a three-stage progressive error correction method, we usually take the first two stages because they are sufficient to eliminate the majority of the errors in many typical NGS datasets. Only in the cases where the data has extensive coverage, such as amplicon sequencing, the option to use the third step is chosen for additional error correction.

### Entropy reduction and information gain after error correction

The error correction effect or the noise/uncertainty reduction by an error correction method in a dataset can be measured by Shannon’s entropy and information gain. For a read dataset *R*, its Shannon entropy *H* is given by

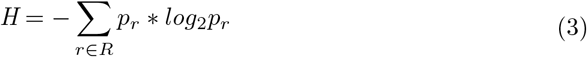

where, *p*_*r*_ is the percentage frequency of *r* in *R*.

An ideal correction should eliminate all the errors/noises in the dataset while not introducing any new errors, or new sequences. Therefore, the entropy *H*^*′*^(*R*^*′*^) of a corrected read dataset *R*^*′*^ should consist of two parts: one is about the original reads, the other is about the wrongly introduced reads. We define *H*^*′*^(*R*^*′*^) as

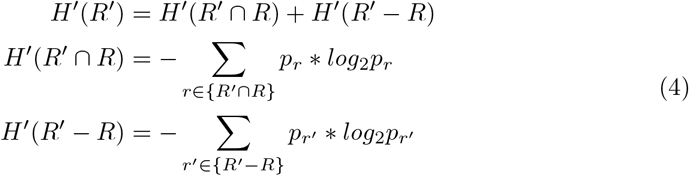

Information gain reflects the amount of information gained from the original state of the reads after the error correction, defined as

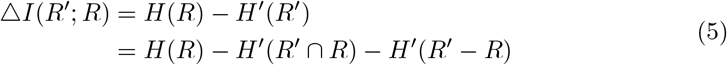

The fewer unique reads is in a dataset, the less uncertainty, and the entropy will decrease after mistaken bases are corrected. To see the error correction effect on the data quality improvement, we propose to visualise the information change via a heatmap of taking items of △*I* as minor rectangular points and marking the wrongly introduced sequences as noises red dots. A visualization of △*I* for *D*1 is shown in Fig. 5 and those for *D*2 - *D*8 are presented at S4-S10 Figs. (details of these datasets are described at the section of Evaluation Criteria of Methods). The primary colour of the heatmaps close to zero strongly suggests that the correction conserves the original high-frequency information by all the methods. The negative and positive scores on the colour bar describe the information gain and loss, respectively. Seen from Fig. 5, noise2read is better than the other methods to reduce noise level as there is nearly no score bigger than zero. Those red points depict information loss brought by wrongly introduced new sequences, leading to new errors to increase false positives and negatives. noise2read does not yield any non-existing reads, and the colour in Fig. 5b darker than that in Fig. 5a implies that noise2read has more information gained from the additional amplicon sequencing correction.

**Fig 5.**
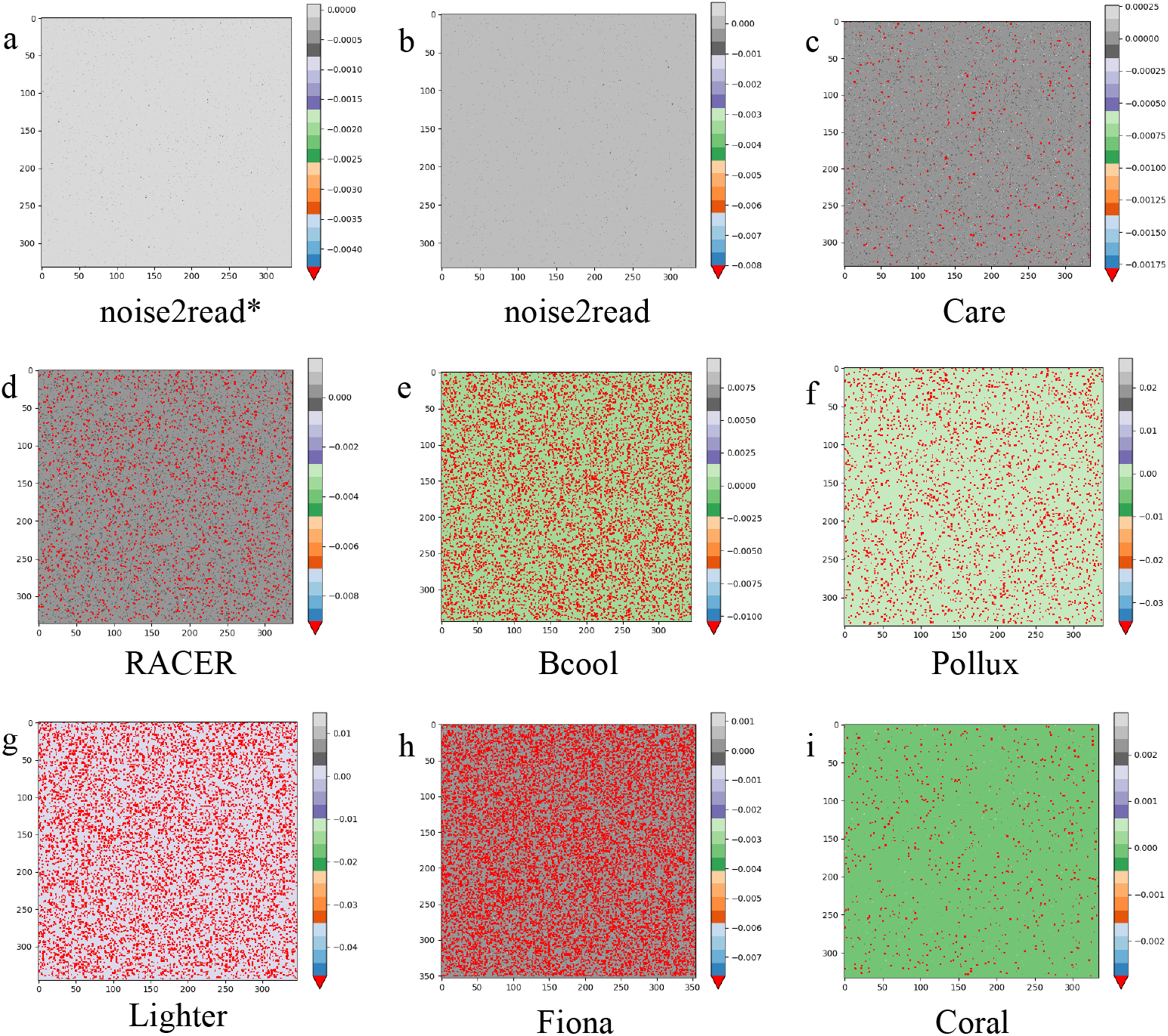
Visualisation of information gain for noise2read*, noise2read, Care [23, 24], RACER [14], Bcool [26], Pollux [17], Lighter [15], Fiona [21], and Coral [20] on dataset *D*1. Here, noise2read* denotes the result without further amplicon correction. Heatmaps **(c, d, e, f, g, h)** and **(i)** display the incorrectly introduced reads as red points. The number of red points shown on each heatmap corresponds to 502, 2310, 7808, 2935, 8523, 13899 and 722, respectively.

Moreover, to intuitively quantify the information gain or loss, we considered the changes only in low-frequency sequences before and after error correction. We denote the frequent reads as a subset *FR*_*τ*_ of *R*, and we calculate the entropy by removing *FR*_*τ*_ from *R* or *R*^*′*^. Then, we focus on the entropy change given by

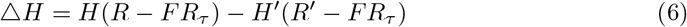

where *τ* is a threshold for defining high-frequency reads, *H*(*R − FR*_*τ*_) and *H*^*′*^(*R*^*′*^ *− FR*_*τ*_) represent the non-frequent reads’ entropy before and after correction, respectively.

We calculated the entropy change △*H* for the sequencing datasets *D*1 - *D*8 and five simulated datasets *D*9 - *D*13 after error correction (details shown in Table 1). noise2read achieves the most considerable information gain on all these datasets. Specifically, the increased information by noise2read on the simulation datasets outperforms the other methods. The extensive information gain is because noise2read can rectify almost all the errors in the simulated datasets.

**Table 1.** Non-frequent reads’ information gain (△*H* with *τ* = 4) on the UMI-based wet-lab datasets *D*1-*D*8 and simulated datasets *D*9-*D*13. Best scores are highlighted in bold.

| Datasets | Methods |  |  |  |  |  |  |  |
| --- | --- | --- | --- | --- | --- | --- | --- | --- |
|  | Coral | RACER | Fiona | Lighter | Pollux | Bcool | Care | noise2read |
| D1 | 0.588 | 6.023 | 4.077 | 4.686 | 2.508 | 0.916 | 1.776 | **6.409** |
| D2 | 0.770 | 6.116 | 4.393 | 4.842 | 2.534 | 0.023 | 1.557 | **6.339** |
| D3 | 0.921 | **6.369** | 4.891 | 5.505 | 3.044 | 2.333 | 0.469 | 6.239 |
| D4 | 0.590 | **6.215** | 4.638 | 5.083 | 3.011 | 1.702 | 0.907 | 6.202 |
| D5 | 0.752 | **6.360** | 4.589 | 5.127 | 2.821 | 0.046 | 1.302 | 6.297 |
| D6 | 0.970 | 6.316 | 4.800 | 5.026 | 2.930 | 1.986 | 1.357 | **6.567** |
| D7 | 0.898 | **6.306** | 4.883 | 5.422 | 3.096 | 0.100 | 0.446 | 6.194 |
| D8 | 1.156 | 5.974 | 4.344 | 4.604 | 3.014 | 1.824 | 1.598 | **6.187** |
| Average | 0.831 | 6.210 | 4.577 | 5.037 | 2.870 | 1.116 | 1.176 | **6.304** |
| D9 | 0.621 | -1.349 | -0.209 | 1.362 | 0.462 | 1.874 | 2.941 | **10.484** |
| D10 | 2.072 | -0.673 | 3.318 | 3.040 | 3.152 | 0.960 | 1.721 | **12.733** |
| D11 | 2.849 | 0.273 | 2.129 | 3.910 | 3.459 | 0.723 | 1.483 | **12.380** |
| D12 | 1.250 | 0.039 | 2.476 | 4.535 | 2.784 | 1.103 | 0.881 | **12.249** |
| D13 | 1.159 | 0.811 | 2.991 | 5.193 | 3.159 | 1.725 | 0.909 | **13.161** |
| Average | 1.590 | -0.180 | 2.141 | 3.608 | 2.603 | 1.277 | 1.587 | **12.201** |

### Performance comparison on UMI-contained sequencing datasets

The above eight datasets *D*1-*D*8 are subsets of wet-lab data sequenced using a UMI-contained high-fidelity sequencing technique (a.k.a. safe-SeqS) [29, 30]. Detailed generation of these datasets can be found in the first subsection of Evaluation criteria at the section Methods. We evaluated the performance of noise2read in comparison with seven other computational error correction methods at both the base-level and read-level under various metrics, including recall (TPR), TNR, fall-out (FPR), FNR, Area Difference (AD), Precision, Positive Gain, Accuracy, and Purity Entropy (E). Detailed definitions of these metrics can be found in Metrics section of Methods.

The comparative performance with the seven methods Coral [20], RACER [14], Fiona [21], Lighter [15], Pollux [17], Bcool [26], and Care [23, 24] are summarized in Fig. 6a for *D*1 and those in S11 Fig and S12 Fig for *D*2-*D*8. S2-S9 Tables. are provided to further supplement the results. Our method noise2read has achieved the best performance on all the datasets under all the metrics.

**Fig 6.**
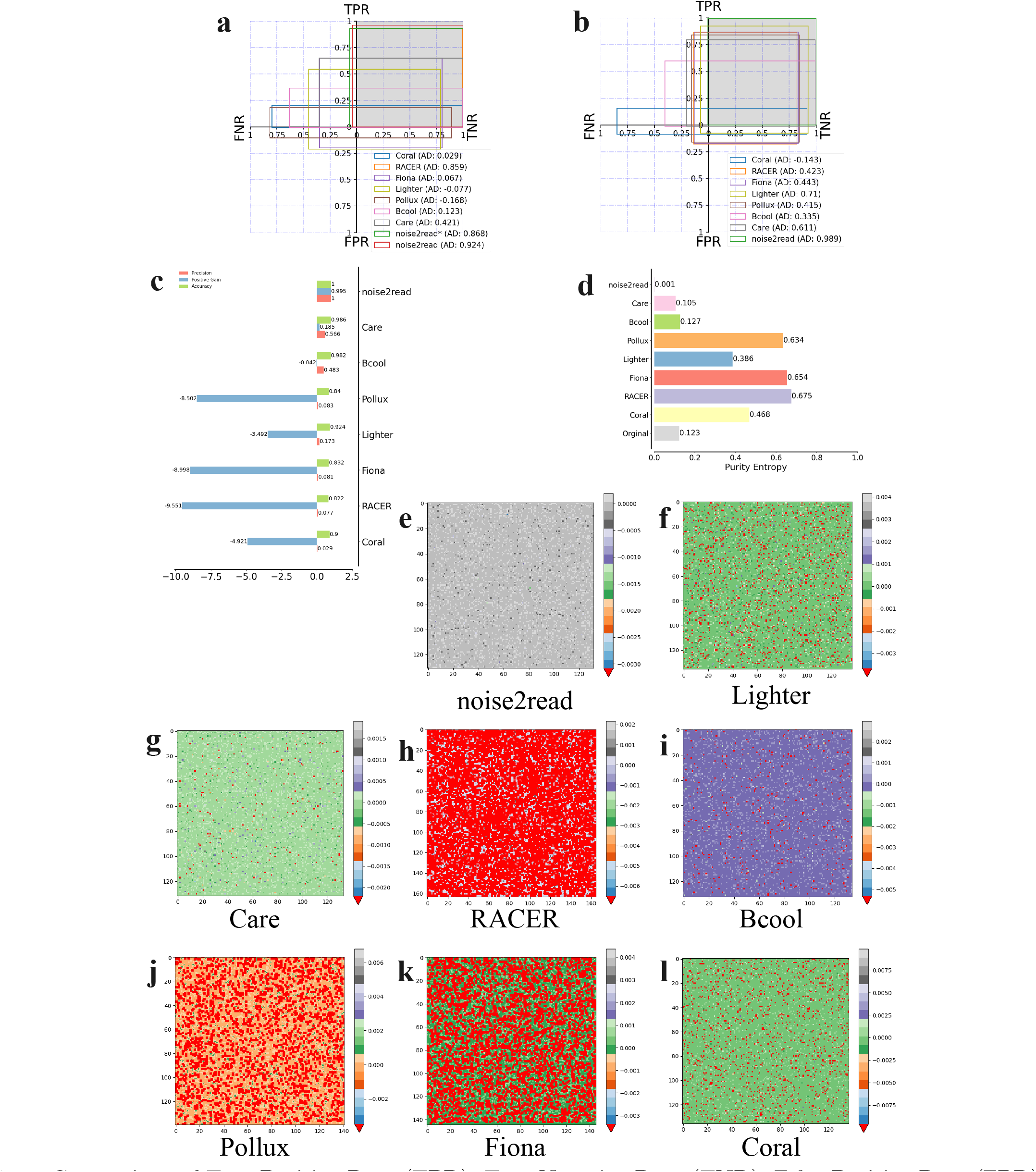
Comparison of True Positive Rate (TPR), True Negative Rate (TNR), False Positive Rate (FPR), False Negative Rate (FNR) and Area Difference (AD) at the read-level for noise2read and seven other methods on the UMI-based wet-lab dataset *D*1 is shown in (a). noise2read* denotes the result without amplicon correction. Performance comparisons at the read-level on simulated dataset *D*9 are shown in (b-d). Information gain visualisations for *D*9 are presented in (e-l). Heatmaps in (f, g, h, i, j, k, l) display the red noise, with 1223, 164, 9698, 377, 2378, 3651 and 1255 points, respectively.

The high-TPR and low-FNR performance indicate that noise2read can turn most noise while leaving the lowest number of actual noise as signals; The high TNR illustrates that noise2read can introduce fewer new errors by preserving most signals unchanged, while the low FPR suggests that noise2read introduces none or few new noises without bringing up any non-existing sequences after the correction process.

In detail, noise2read surpassed all the other methods on *D*1, achieving a score 0.924 higher than the second-best method RACER which has a score of 0.859. Notably, noise2read exhibited exceptional performance in Recall, Precision, Positive Gain, Accuracy, and Purity Entropy, as evidenced by the values in Table 2. The positive gain percentage of noise2read is 7.26% and 48.15% higher than RACER and Care. noise2read and its amplicon mode achieved the finest purity entropy of 0.05 and 0.077, sounder than the second-best method RACER which has a score of 0.110.

**Table 2.** **TP, FP, FN, TN, Recall, Precision, Positive Gain, Accuracy and purity Entropy(E) at the read level for comparing our method noise2read (decomposed at different stages) with seven existing methods on the UMI-based wet-lab dataset *D*1. Best scores are highlighted in bold**.

|  |  | Metrics |  |  |  |  |  |  |  |  |
| --- | --- | --- | --- | --- | --- | --- | --- | --- | --- | --- |
| Methods |  | TP | FP | FN | TN | Recall | Precision | Positive Gain | Accuracy | E |
| noise2read | stage 1 | 81630 | 0 | 30391 | 693059 | 0.729 | 1.000 | 0.729 | 0.962 | 0.232 |
|  | stage 2 | 104373 | 0 | 7648 | **693059** | 0.932 | **1.000** | 0.932 | 0.991 | 0.077 |
|  | stage 3 | **107717** | 201 | **4304** | 692858 | **0.962** | 0.998 | **0.960** | **0.994** | **0.050** |
| Coral |  | 22677 | 4249 | 89344 | 688810 | 0.202 | 0.842 | 0.165 | 0.884 | 0.518 |
| RACER |  | 104589 | 4347 | 7432 | 688712 | 0.934 | 0.960 | 0.895 | 0.985 | 0.110 |
| Fiona |  | 72792 | 136472 | 39229 | 556587 | 0.650 | 0.348 | -0.568 | 0.782 | 0.757 |
| Lighter |  | 61330 | 145999 | 50691 | 547060 | 0.547 | 0.296 | -0.756 | 0.756 | 0.802 |
| Pollux |  | 20537 | 73710 | 91484 | 619349 | 0.183 | 0.218 | -0.475 | 0.795 | 0.732 |
| Bcool |  | 40970 | 3889 | 71051 | 689170 | 0.366 | 0.913 | 0.331 | 0.907 | 0.447 |
| Care |  | 72673 | 52 | 39348 | 693007 | 0.649 | 0.999 | 0.648 | 0.951 | 0.3 |

The progressive process gradually converts noise into signals; for example in Table 2, the 1st, 2nd and 3rd stages convert 72.9% (81,630), 93.2% (104,373), and 96.2% (107,717) of the errors into signals on *D*1, respectively. noise2read is mainly designed for any short-read sequencing data whenever PCR is involved. Without the 3rd step, it also achieves sound performance (refer to S2-S9 Tables.) by restoring most erroneous reads into their normal states and not introducing false positive reads. The 3rd step for further correction on amplicon sequencing data maintains fewer original error-free reads than the second stage and correspondingly introduces more noise but not new sequences. RACER can rectify 93.4% of the noise but newly introduces almost 22 times the number of new errors compared to our method. Care newly introduces 52 false positives but can only correct 64.9% of the erroneous reads. The other methods can only correct less than 65% of the errors but simultaneously give rise to thousands of new mistakes.

We conducted additional performance comparisons between our noise2read and ten state-of-the-art approaches on eight UMI-based ground-truth datasets all established previously by the literature [30]. We refer these datasets as D34 to D41. Detailed information about these datasets can be found at Supplemental Table 1.

To conduct a fair comparison, we employed the evaluation procedures presented in the literature [30] to compute the confusion matrix at the read level. By this definition, a read is deemed erroneous if even a single base is incorrect, namely, a read is error-free only when all its bases are correct.

Table 3 shows the comparative performance of noise2read on D41 in comparison with Bless [18], Coral [20], Lighter [15], Reckoner [19], Sga [25], BFC [16], Pollux [17], Fiona [21], RACER [14], and Care [23, 24]. For comparative performance on D34-40, please see S17 Table. and S18 Table.. These comparison results highlight that noise2read consistently achieved the highest number of true positives on all of D34-D41, except for Fiona’s TP on D41, which is slightly bigger than noise2read. Importantly, noise2read demonstrates the lowest count of false positives among all these datasets. noise2read performs exceptionally good in Precision, Accuracy, AD and Positive Gain on all of the eight benchmark datasets.

**Table 3.**
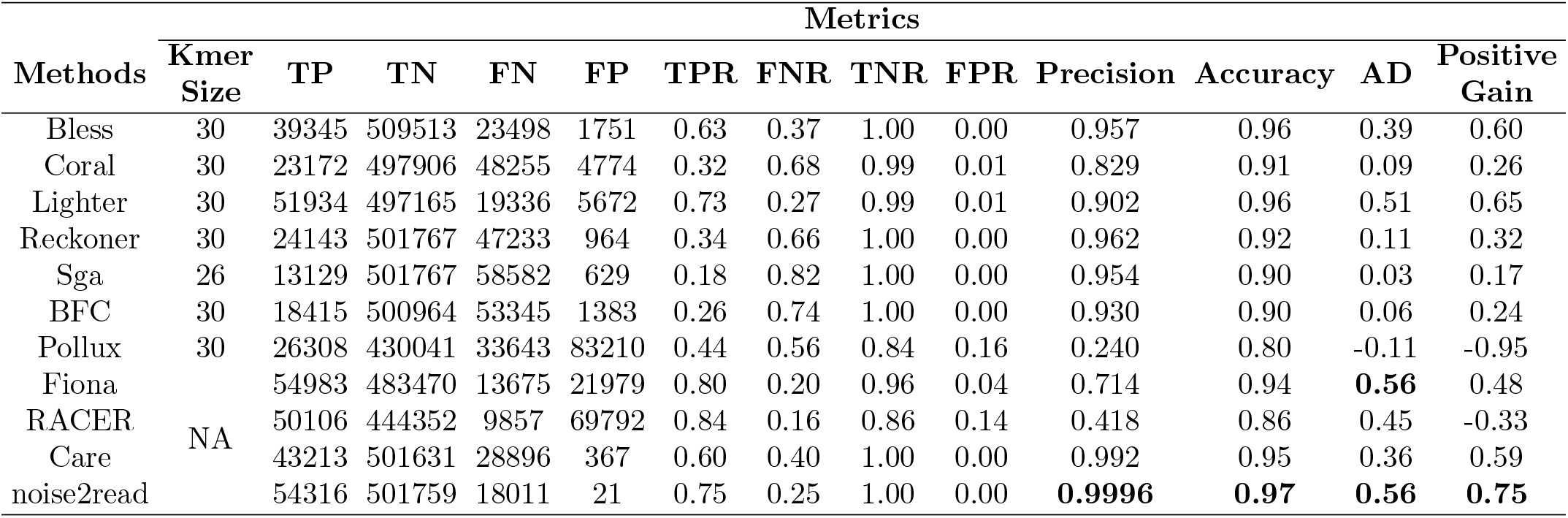
The comparative performance between noise2read and methods Bless [18], Coral [20], Lighter [15], Reckoner [19], Sga [25], BFC [16], Pollux [17], Fiona [21], RACER [14], and Care [23, 24] on the UMI-based benchmark dataset *D*41 previously established in the literature [30].

### Performance on simulated short-read datasets and those with artificially modified bases

Based on actual sequencing datasets of various read lengths of 75bp, 101bp and 150bp, we simulated datasets *D*9-*D*13 with mimic UMIs. *D*9 was simulated from a single-end dataset with SRA accession SRR12060401, while *D*10 and *D*11 were simulated using the forward sequencing data (R1) and the reverse complementing sequence data (R2) from a paired-end data with SRA accession SRR9077111. Similarly, *D*12 and *D*13 were simulated using R1 and R2 from a paired-end data with SRA accession SRR11092062. The simulation procedure is presented in Evaluation Criteria of Methods.

Error correction performance are shown in Fig. 6b-l, S13 Fig and S10 Table. for dataset *D*9 and S14-S21 Figs. and S11-S14 Tables. for datasets *D*10-*D*13. noise2read super outperforms the other methods under all the metrics on all the simulated datasets. Specifically for the AD performance, noise2read (0.989) has 39.3%, 61.9% and 123.3% higher performance than that of Lighter(0.71), Care(0.611) and Fiona (0.443) (Fig. 6b). noise2read reaches the best precision, gain and accuracy and achieves a substantial positive gain (Fig. 6c).As shown in Fig. 6d, noise2read is the only method significantly decreasing the purity Entropy after correction. Information gain visualisations in Fig. 6e-l indicate the information is still dominated by most of the original signal after correction. All the other methods wrongly introduced new sequences (in a number of 164 to 9698) after correction. The other methods’ performance fluctuates widely. For instance, at the read level, the performance ranking of the top three methods in terms of AD on dataset *D*9 (Fig. 6b) is Lighter, Care and Fiona. However, the performance ranking is Fiona, Care, and Coral on *D*10 (S14(b) Fig), and Lighter, Fiona and Care (S16(b)Fig) on *D*12.

Moreover, we simulated four single-end miRNA datasets (denoted by *D*14-*D*17) using the simulation procedure proposed in miREC [27] plus an additional step of mimicking UMIs to these datasets. Of them, *D*14-*D*15 contain substitution and indel errors, while *D*16-*D*17 contain only substitution errors. Comparison results between noise2read (with or without high-frequency ambiguous error prediction) and miREC are shown in Table 4 and S15 Table. noise2read can rectify more errors than miREC, achieving more TP and less FN after correction. The miREC method and noise2read can achieve similar, reasonably good results in accuracy, precision and fall-out (S15 Table). However, from the recall, Positive Gain, purity Entropy *E* and information gain △*H* performance on all four datasets, noise2read is better than miREC.

**Table 4.** Performance comparison between noise2read (with and without enabling the high-frequency ambiguous error prediction) and miREC on miRNA simulation datasets *D*14-*D*17 at the read level.

| Datasets | Methods | Metrics |  |  |  |  |  |  |  |
| --- | --- | --- | --- | --- | --- | --- | --- | --- | --- |
| | | TP | FP | FN | TN | Recall | Positive Gain | $\Delta E$ | $\Delta H$ |
| D14 | miREC | 4224 | 0 | 267 | 218980 | 0.941 | 0.941 | 0.129 | 4.755 |
|  | noise2read | 4411 | 11 | 80 | 218969 | 0.982 | 0.980 | 0.137 | 9.053 |
|  | noise2read<sup>a</sup> | 4409 | 3 | 82 | 218977 | 0.982 | 0.981 | 0.137 | 9.053 |
| D15 | miREC | 4135 | 5 | 271 | 219060 | 0.938 | 0.937 | 0.126 | 4.821 |
|  | noise2read | 4305 | 7 | 101 | 219058 | 0.977 | 0.975 | 0.134 | 8.485 |
|  | noise2read<sup>a</sup> | 4308 | 4 | 98 | 219061 | 0.978 | 0.977 | 0.134 | 8.485 |
| D16 | miREC | 6418 | 16 | 309 | 216728 | 0.954 | 0.952 | 0.179 | 5.301 |
|  | noise2read | 6577 | 17 | 150 | 216727 | 0.978 | 0.975 | 0.186 | 8.184 |
|  | noise2read<sup>a</sup> | 6579 | 16 | 148 | 216728 | 0.978 | 0.976 | 0.186 | 8.184 |
| D17 | miREC | 6398 | 0 | 306 | 216767 | 0.954 | 0.954 | 0.179 | 5.337 |
|  | noise2read | 6577 | 1 | 127 | 216766 | 0.981 | 0.981 | 0.187 | 8.769 |
|  | noise2read<sup>a</sup> | 6574 | 0 | 130 | 216767 | 0.981 | 0.981 | 0.187 | 8.769 |

<sup>a</sup> Performance by noise2read without prediction of errors between high-frequency reads.
<sup>b</sup> High-frequency threshold $\tau = 4$ .

### Error correction changes the abundance level by 12% for the reference genome of SAS-Cov-2 and by 52.12% for Monkeypox

The SARS-CoV-2 and Monkeypox viruses have severely affected the health of human beings. The reference genome sequences have been extensively utilized to understand the origin and phylogeny, and also as a fundamental framework for the design of mRNA vaccines. We investigate how much abundance is changed for the reference genomes of these two viruses after our algorithm noise2read rectifies the base errors contained in the short-read sequencing data of the reference genomes. The study will help understand the within-host viral mutants of the reference genome and their abundance compositions.

We removed human-related reads from paired-end sequencing dataset SRR11092062 [31] using Bowtie2 [32] and the human genome reference GRCh38.p13. The paired R1 and R2 of SRR11092062 after the filtering process were denoted as *D*18 and *D*19, respectively. Then, we performed read alignments for all the reads in *D*18 and *D*19 using Bowtie2, and extracted only those reads that have perfect matches with the viral reference genome MN996528.1 [31] to calculate the base coverage (abundance level) of the viral reference genome. We note that the reference genome MN996528.1 of SARS-CoV-2 was de novo assembled from paired-end short-read sequencing dataset SRR11092062 after human-related reads were filtered.

Our method noise2read corrected 181,360 erroneous reads in *D*18 and corrected 138,575 erroneous reads in *D*19. In particular, 2328 corrected reads can be perfectly aligned to the genome. This leads to the base coverage of the reference genome increased by 12% on average, namely an elevation of the sequencing coverage from 96.07 to 107.75 (or from the 19,144 perfectly matched reads to 21472) (Figure 7a, b). Figure 7c depicts a frequency distribution under a scaled density curve for the coverage difference after error correction. Some regions of the genome maintain the same low level of base coverage without change after the reads correction. Interestingly, some base coverage becomes lower after the error correction (e.g., position 13541) suggesting there exists a within-host mutant that gains new support of perfectly matched reads turned from the reference genome.

**Fig 7.**
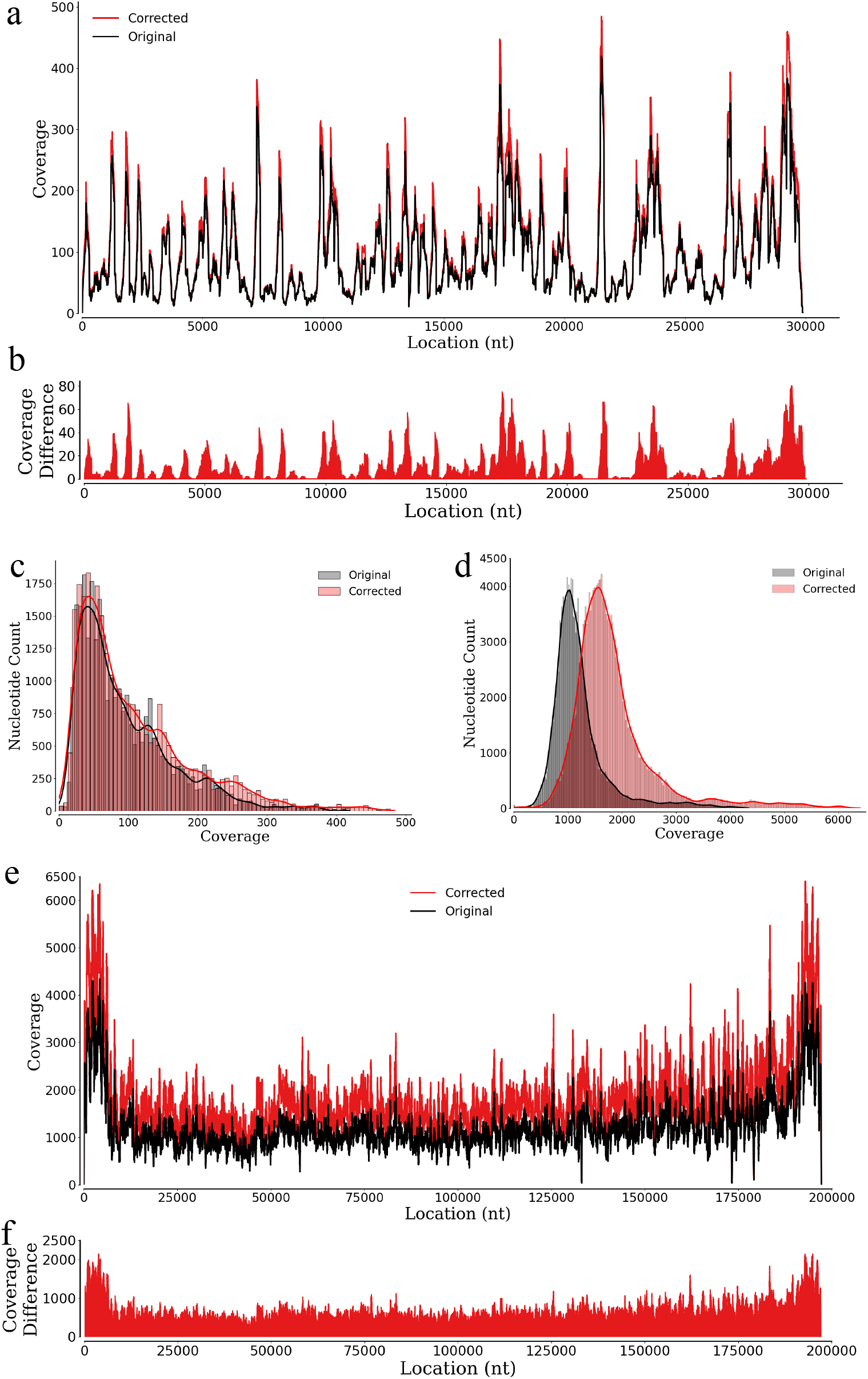
Comparison of base coverage before and after correction for SARS-Cov-2 and Monkeypox virus genomes using perfectly matched reads. (a) and (b) show the base coverage and its difference for the SARS-Cov-2 data, while (e) and (f) present the base coverage and coverage difference for the Monkeypox virus data. Frequency distributions of the coverage difference are shown in (c) and (d) with scaled density curves for the SARS-Cov-2 and Monkeypox virus data, respectively.

In the abundance change analysis for the Monkeypox virus, we used the paired-end whole-genome sequencing dataset SRR22085311 (its paired R1 and R2 denoted as *D*20 and *D*21 here) and the reference genome GCA 025947495.1 [33]. Our noise2read rectified a huge number of erroneous reads in *D*20 (400,622 out of 3,599,812 reads, i.e., 11.13%), and another huge number of erroneous reads in *D*21 (456,242 out of 3,599,812 reads, i.e., 12.67%). Figure 7e presents a coverage comparison chart before and after the error correction, alongside a coverage difference chart (Figure 7f), where it can be seen that the average base coverage of the reference genome is increased by 52.12% from depth 1216.75 to 1850.95 after a huge number of 651,410 reads were retrieved to perfectly align with the genome.The frequency distribution of the base coverage differences as another angle viewing the abundance change for the Monkeypox virus is presented in (Figure 7d), where the abundant and perfectly matched reads aligned to the genome are highlighted again. Especially, those positive shifts towards a higher coverage (Figure 7e,f) confirm much more about the ground truth of the known reference genome and the detection of possible new variants of the genome.

### Accurate error correction improves detection of isomiRs and refines SNPs profiling

MicroRNAs (miRNAs), non-coding RNA molecules approximately 22 nucleotides long, can modulate gene expression post-transcriptionally through the silencing and decay of target mRNAs [34]. Dysregulation of miRNAs plays crucial roles in many biological mechanisms, and it is also a main reason in cancer and autoimmune disorders [35, 36]. By miRNA sequencing, various types of isoforms (i.e., isomiRs) have been detected [37]. However, whether the base differences found in the isomiRs are actual biological variations or synthetic artefacts due to the PCR or sequencing errors or both is difficult to judge. Here, we study how our error correction changes the identification and quantification of isomiRs from short RNA-seq datasets and how it refines the profiling of known single-nucleotide polymorphisms (SNPs) in isomiRs.

We downloaded ten single-end small RNA-sequencing datasets of lymphoblastoid cell lines from five population groups in the 1000 Genomes Project [38]. These datasets (denoted as *D*22-*D*31 here) were cleaned by removing the adapter sequences via cutadapt [39]. We used IsoMiRmap [40] under the setting of pre-defined miRNA reference sets from the database miRBase(v22) [41] as a “miR-space” to quantify known isomiRs and SNPs for *D*22-*D*31 before and after our sequencing error correction. IsoMiRmap tags an identified isomiR as an exclusive isomiR if it only exists in the miR-space with one or more occurrences but not elsewhere in the human reference genome, otherwise recognized as an ambiguous isomiR.

These quantification results are summarised in Table 5. The number of unique ambiguous isomiRs is decreased by 24.12%-31.75% or in numbers from 151 to 245, but their total counts are increased by a number between 160 and 640 among the ten datasets after the error correction; the number of exclusive isomiRs is decreased by 34.46%-37.48% but their total counts are increased by a number between 5095 and 14,441. These results suggest that some previously identified isomiRs are artefacts containing sequencing errors rather than natural isoforms. On the other hand for the profiling of the known SNPs, the number of unique SNPs is decreased by 34.13% - 59.09%, and their counts are also decreased by 4.40% - 35.56% except for two increased by 1.41% and 1.61% respectively. This observation unveils that some of the previously annotated SNPs are actually sequencing errors. Similar quantitative and qualitative changes observed in the profiling of these known SNPs in the isomiRs distinguishing true SNPs from sequencing errors enables more accurate annotation of SNPs. The significant change of the isomiRs quantification after correction is because an average of 235,146 (2.62%) sequences were corrected by noise2read in the ten datasets (Table 5).

**Table 5.**
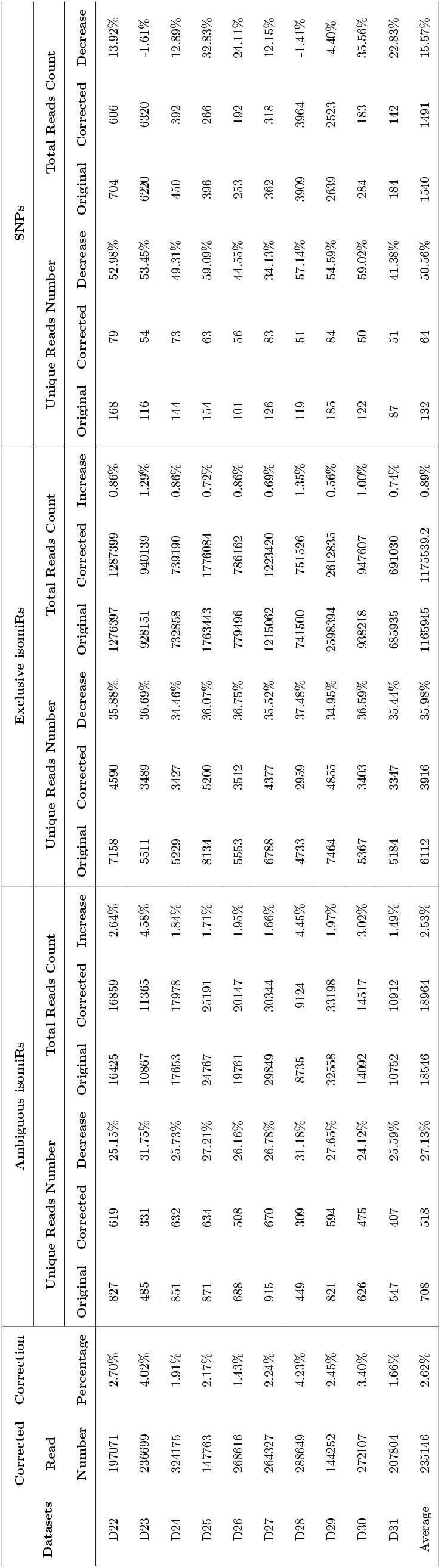
Known isomiRs and SNPs profiling change from miRNA sequencing data before and after correction.

To understand more about the frequency change of isomiRs and SNPs, we categorised the isomiRs according to their original miRNAs, then we utilised scatter graphs with Kepler plots to understand the associations between the number of identical isomiRs and total isomiRs’ count (log_10_ transformation) before and after the error correction of the sequencing reads. The leftward shift on the x-axis (Fig. 8a and 8b for exclusive isomiRs of *D*22 and *D*23,respectively, Fig. 8c, d for ambiguous isomiRs and known SNPs of *D*22, and S22-S24 Figs for the other miRNA datasets) indicates a reduction of the count of unique isomiRs, while the upward change on the y-axis indicates an increase in authentic isomiRs. These significant changes in isomiRs and SNPs highlight the importance of correction for accurately characterizing isomiR and SNP profiles, making contributions to the annotation of isomiRnome.

**Fig 8.**
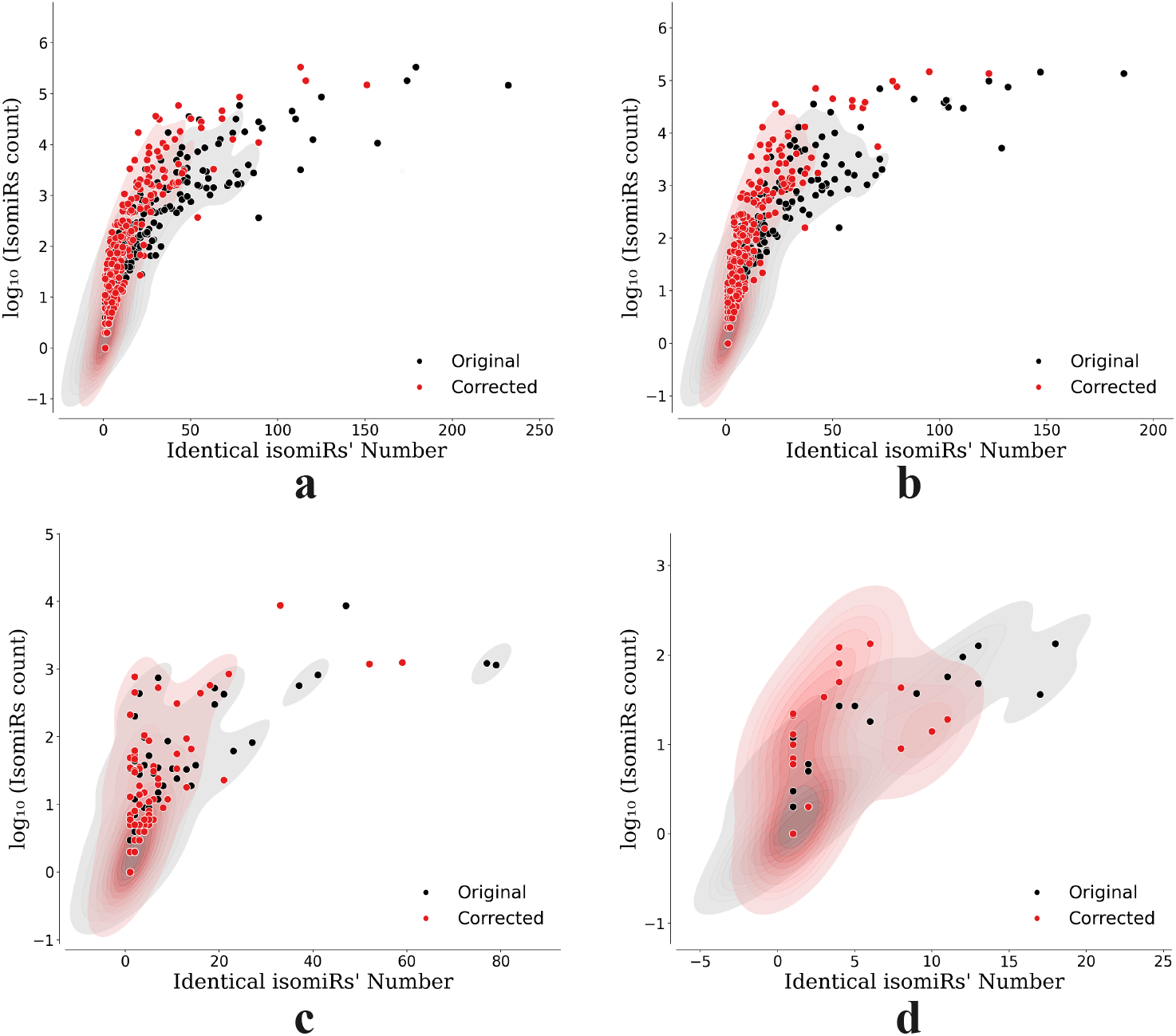
Scatter graphs compare the numbers of isomiRs and known SNPs before and after correction, while the Kepler plots provide additional visualizations of the data distribution. a and b compare the numbers of exclusive isomiRs from *D*22 and *D*23, respectively, and c and d compare the numbers of ambiguous isomiRs and known SNPs from *D*22.

### Accurate error correction significantly improves the quality of outcome sequences edited from a disease gene by ABEs and CBEs

Base editing is a new genome editing technique that uses CRISPR systems and enzymes to introduce point mutations into cellular DNA or RNA for modeling and understanding genetic diseases [42, 43]. However, deciding whether a nucleotide position is exactly editable in a genomic context is inefficient by wet-lab experiments, and the base editors may yield many unexpected genotypic output sequences when the editable window covers multiple target nucleotides. Deep-learning-based prediction tools have been developed to predict the base-editing efficiency and outcome-sequence copy numbers from Adenine and cytosine base editors (ABEs and CBEs) [44]. The training data used by these prediction tools are extracted from short-read DNA/RNA sequencing data. Here, we investigate how much the number of unique reads (unique outcome sequences) changes after our sequencing error correction.

We removed those records in which the target sequence has only one outcome sequence from the training data of HT ABE Train and HT CBE Train used in the literature [44]. Then, we cleaned them to form two datasets (denoted by *D*32 for ABEs and *D*33 for CBEs here), and applied noise2read to *D*32 and *D*33 separately. As a result, the number of unique outcome sequences in *D*32 is reduced by 2309 from 28892 to 26583 (7.99%), and the number of unique outcome sequences in *D*33 is reduced by 5042 from 27312 to 22270 (18.46%). The number reduction of unique outcome sequences is because some low-frequency reads are not a result of base editing but due to sequencing errors. In total, noise2read recognised 5109 erroneous reads in the ABE dataset and 10271 erroneous reads in the CBE dataset and turned all of them into normal states. This error correction has significantly improved the quality of the training data that would be very helpful for enhancing the prediction of base editing efficiencies.

### Runtime and memory consumption

We compared CPU runtime and peak memory used by noise2read with those by Care [23, 24], RACER [14], Bcool [26], Pollux [17], Lighter [15], Fiona [21], and Coral [20] on data sets *D*1-*D*8. We executed all the programs on an Intel(R) Xeon(R) Gold 6238R CPU clocked at 2.20GHz, leveraging 56 CPU cores for parallel computing. For the model training of noise2read, a single Tesla V100S-PCIE-32GB GPU was employed. To gauge memory usage across all the programs, we used the library psutil [59]. These runtime and memory consumption comparisons are presented in Table 6.

**Table 6.**
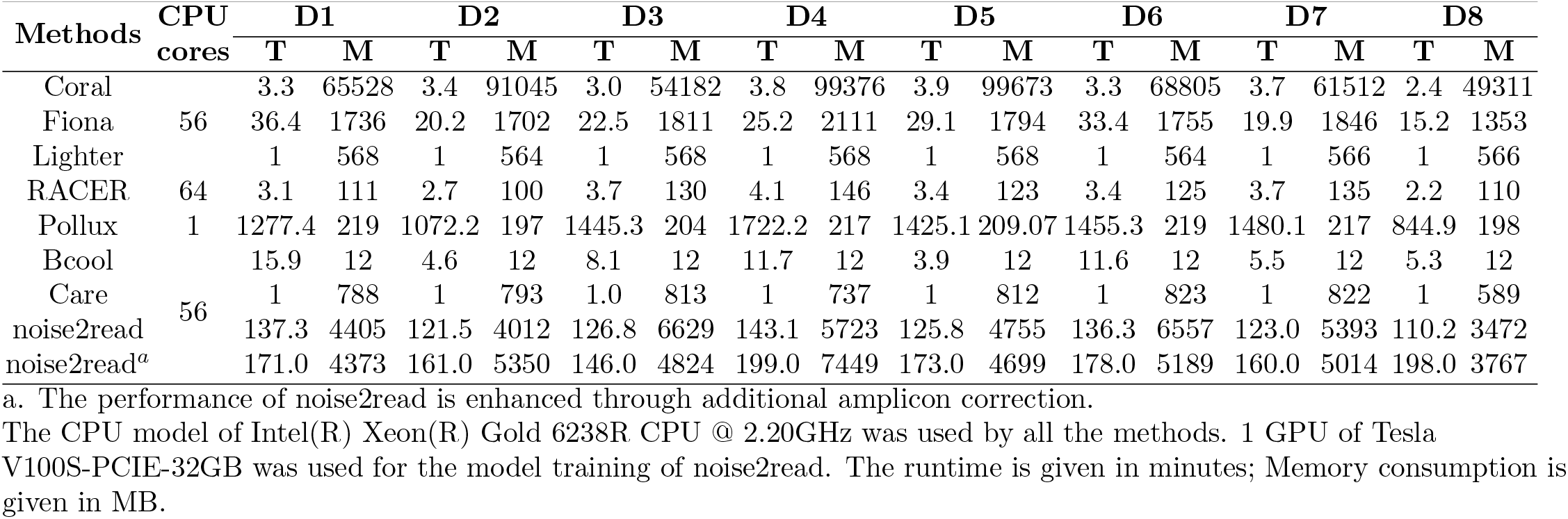
Runtime(T) and Memory(M) usage by different methods on the datasets D1-D8.

Lighter and Care exhibited fast speeds, completing corrections within a minute by taking a small amount of memory consumption for each of the 8 data sets. On the other hand, Pollux had the slowest speed due to its inability to run in parallel. noise2read spent the second highest memory consumption and made the second slowest speed. (We note that we do not suggest using a large number of multiprocessing processes for noise2read to run, as we have observed that those situations could suddenly consume a significant amount of memory, and the program ran out of memory and terminated.)

The separate time consumption by noise2read at its different stages as recorded in the built-in log files across all datasets *D*1-*D*41 are presented in S19 Table. It can be observed that a significant amount of time was spent on tasks such as constructing “2nt-edit-distance read graphs”, “performing feature extraction”, and “model training”. To shorten the running time on large data sets, it is suggested to choose a smaller number of negative samples and set a smaller number of trials (e.g., 20) for the construction of sub-optimal models. Additionally, opting not to predict errors within high-frequency reads will also save noise2read a substantial amount of time and memory usage but fortunately without much performance sacrifice on error correction. We note that although noise2read is slow, it never introduces any non-existing reads into the datasets. This is a unique merit in all of current sequencing error correction methods.

While the speed of error correction is undeniably a crucial factor in evaluating the performance of a correction method, an even more critical consideration is whether the method introduces new errors. A fast error-introducing method damages the quality of the whole dataset and may become unexpectedly harmful to downstream data analysis, although its fast speed is advantageous in the preprocessing error correction stage. Our method does not have this speed advantage so far, but it never introduces new errors, guaranteeing the integrity of the datasets. In future work, we consider efficiency tricks to improve the speed of feature extraction and machine learning.

## Discussion

A long-standing problem in sequencing data analysis is how to reduce sequencing base errors and erroneous reads as much as possible before any downstream applications. Existing short reads correction methods utilize biochemical-based experimental designs such as unique molecular identifiers (UMIs) to count and track molecules [10], or take computational methods including *k*-mer-methods [13–19], multiple sequence alignment based methods [20–24], and the other methods [25, 26]. One limit of the UMI-based strategies is that errors/mutations can also happen at UMIs. Serious concern about the computational methods is that they have significantly overcorrected reads by introducing pseudo new sequences or shifting one type of error into another, often leaving numerous reads uncorrected. Some of these methods only focus on restoring substitution mistakes but do not support indels’ correction. Besides, instance-based methods such as miREC [27] were designed to handle specific sequencing data type miRNA sequencing reads. And it assumes that frequent sequences contain no mistakes, thus it cannot be used to correct potential errors between high-frequency reads or cannot deal with those singletons with no relationships to the high-frequency reads.

Following the principle of the PCR erring incidents and sequencing process, we constructed special graphs of short reads to capture the relationships between edit-erring and error-free reads. Through novel modelling of the errors between high-frequency reads and their high- or low-frequency neighbours as a classification problem, we have successfully predicted almost all the errors using machine learning techniques. Validation experiments on the UMI-based wet lab and simulated datasets of known ground truth have demonstrated that the proposed noise2read algorithm can eliminate most of the PCR and sequencing errors without introducing any non-existing sequences into the read set.

Moreover, we investigated the impact of error-corrected data on downstream data applications. We have found that: (1) The abundance level change of the reference genome of SARS-CoV-2 and Monkeypox after the sequencing error correction is remarkable, which may allow us to rethink how to get a precise genome sequence for these viruses; (2) For the isomiRs and SNPs profiling, the counts of some isomiRs and SNPs are decreased while some others are increased, which is of great significance to identifying actual isomiRs and SNPs and re-annotating the isomiRnome. (3) Both ABE and CBE should have higher base editing efficiency than currently estimated. The accurate and higher base editing efficiency with correct preprocessing may improve the original deep-learning prediction accuracy. Altogether, these observations and advantages lay down strong evidence to question the accuracies of current downstream research outcomes and open new avenues to conduct downstream analysis whenever short-read data are adopted.

A small edit distance such as 1 or 2 is currently used to define the edges of *rG*(*R*). When the edit distance threshold *Emax* is enlarged, more edges will be created for *rG*(*R*) and possibly more erroneous reads will be identified. The tradeoff is that the computational complexity of constructing these new edges is exponential while newly identified erroneous reads become less and less when *Emax* increase. In fact, these erroneous reads constitute an extremely small percentage (*<* 0.16%) of the total erroneous reads in theory. In future work, we will test the computational complexity when *Emax* is set as 3, and explore how to change the correction steps.

The speed and memory usage of noise2read still needs improvement, especially the parts for building the 1nt- and 2nt-edit-distance read graphs and AutoML training for prediction. The easy-usable and automatic tuning of the classifiers’ parameters facilitates wide-range explorations, but we note that noise2read may yield a slightly different result at different trials, even setting the same seeds. We also note that noise2read will derive more false positives when dealing with errors between high-frequency reads of extremely short length (e.g. *<* 30bp). This limit may be overcome by extracting more or fewer features from the reads. Furthermore, we already attempted using deep learning architecture (e.g., CNN, LSTM) to detect the errors, but a better performance was not achieved than by current noise2read. To make noise2read from outstanding to an exceptional tool, novel conception of new features for the short reads and attention-based deep learning models are potential solutions as our future work.

## Methods

### Implementation of our noise2read algorithm

The proposed algorithm noise2read was implemented using the python language and packaged as an easily-usable prototype publicly available. It mainly consists of modules for the construction of 1nt- and 2nt-edit-distance read graphs, for the creation of the training sets edit-erring-READS and error-free-READS for machine learning, for Auto Machine Learning (AutoML) prediction, and for the error correction steps.

### Special considerations in the construction of edit-distance graph of short reads

By setting a high-frequency threshold *τ*, noise2read finds the 1nt- or 2nt-edit-distance edges between unique high-frequency reads (with frequency *> τ*) and all the other unique reads in a read dataset, and then it takes all of these unique reads as nodes, their counts as attributes and the detected associations to build a graph. The rationale for not detecting the 1nt- or 2nt-edit-distance read pairs in the low-frequency reads is that it is computationally challenging and meaningless to distinguish whether one read of low abundance is mutated from the other low-frequency read (e.g., it is hard to determine if there are mutations or sequencing errors between two reads each with a frequency of one and with one base or two-base difference). The rationale for 2nt-edit-distance error correction is that some NGS data contains two base errors in some long read (e.g., 150 bp), and we set a threshold *l* (e.g., 30) of the sequence’s minimum length to determine whether to perform 2nt-edit-distance error correction.

Noise2read does not perform a pairwise alignment for searching the 1nt- or 2nt-edit-distance edges between the high-frequency reads and all the other reads in the read set. Instead, it enumerates all the possible 1nt- or 2nt-edit-distance (substitutions only for the 2nt) reads for all of the high-frequency reads and stores them in the Python Set. Then, it invokes the Python built-in function intersection to obtain the edges. It may not be the best way to find all the edges using hash tables in this manner. However, such a strategy can find all required edges instead of finding an approximate number of edges.

We constructed the 2nt-edit-distance-based graph by searching only substitution relations as edges. This idea is based on the observation that substitutions are the most prevalent type of sequencing error [45], and on that ambiguous nucleotides are often denoted by the symbol ‘N’ [46, 47] during sequencing. Moreover, NGS read lengths are usually consistent and fixed in a single sequencing run, owing to the fixed number of sequencing cycles in technologies like Illumina sequencing. This uniform read length is achieved since the read size is directly tied to the number of sequencing cycles performed, and each cycle corresponds to the sequencing of a single base. On the other hand, if a deletion or insertion exists in the read, the sequence length will change, and such a sequence will not appear in a uniform-length sequencing dataset. Noteworthy, noise2read can handle indel errors when insertion or deletions are represented by the symbol ‘N’.

### Construction of edit-erring-READS and error-free-READS as training data

By defining a maximum frequency threshold *τ*_*err*_ (*τ*_*err*_ ≤ *τ*), we considered two kinds of erroneous reads: genuine errors and ambiguous errors. Genuine errors are referred to those leaf nodes whose frequency *τ* ^*′*^ is less than or equal to *τ*_*err*_ (*τ* ^*′*^ ≤ *τ*_*err*_) and which have a neighbouring node with a higher frequency than *τ*. This set of erroneous reads is denoted as edit-erring-READS. Genuine errors can be directly rectified to their correct states. While we define two kinds of ambiguous errors: **(a)** those nodes (reads) *r* with a low-frequency *τ* ^*′*^ that are each connected to multiple (≥ 2) high-frequency nodes; **(b)** wrongly sequenced reads existing between a pair of similar high-frequency reads as the second kind of ambiguous error instances. In other words, in the constructed 1nt-edit-distance-based read graph, if there are edges between two similar high-frequency sequences, there may be sequencing errors between them. Moreover, amplicon sequencing utilises ultra-deep PCR amplifications for a specific gene target and supports hundreds to thousands of amplicons multiplexed sequencing in one assay to achieve high coverage, but ultra-deep PCR simultaneously amplifies PCR errors. To this end, we further construct a 1nt-edit-distance-based read graph for amplicon sequencing data and consider those reads of frequencies less than 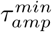 (e.g., 50) as potential amplicon errors mutated from its neighbouring reads of extremely high-frequency larger than 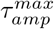 (e.g., 1500).

We consider isolated nodes of high frequencies bigger than *τ* as error-free reads. We take those isolated nodes of high frequencies in the 1nt- or 2nt-edit-distance graphs to build the training set error-free-READS.

### Auto Machine Learning (AutoML) prediction

Unlike the direct rectification of genuine errors into their original state, we model whether a high-frequency read contains true mutations or sequencing errors from its high- or low-frequency neighbours as a classification problem. We created an auto Machine Learning (AutoML) module for its end-to-end prediction. The flowchart illustrated in Fig. 9 outlines the steps involved in the AutoML module.

**Fig 9.**
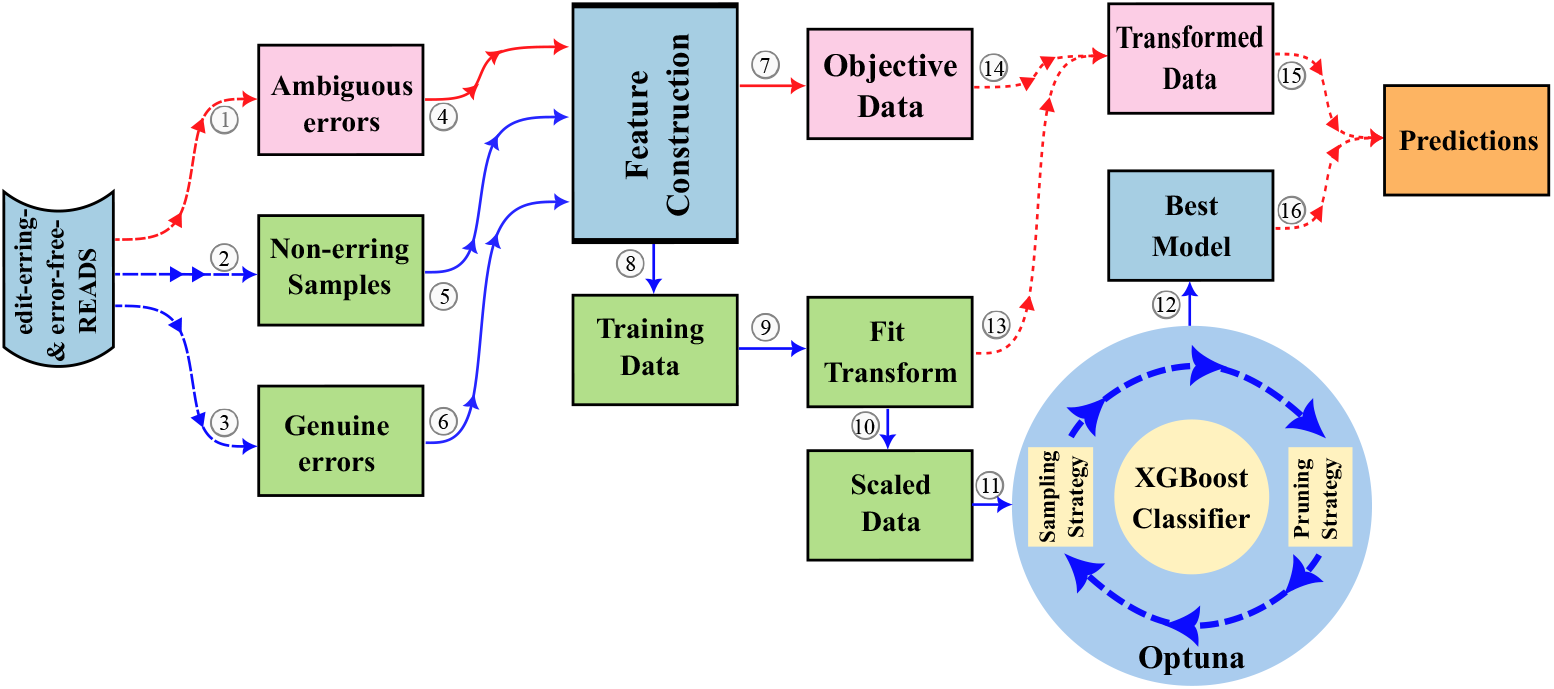
An overview workflow of the Auto Machine Learning (AutoML) module for end-to-end prediction.

### Formulation of the classification problem

We consider edit-erring-READS as positive training instances, while error-free-READS as negatives. For a low-frequency node with a degree greater than two, we calculate its probability of mutation from all its high-frequency neighbouring nodes and take the node with the highest probability as its correct sequence. For the second type of ambiguous error prediction, we integrate the predicted results of the first kind into the training data. In the current version, we only use the predicted ambiguous samples as negative samples for high-ambiguous error prediction to reduce training time and complexity. The mutation observed in high-frequency reads exhibits a bidirectional nature. Therefore, we only consider the prediction result with a higher probability when the bidirectional predictions match. In other words, if the absolute difference between the probabilities of the two-way predictions is less than a specific value, we discard the prediction; otherwise, we choose the prediction having a higher probability.

### Feature representation for the training and objective data

A short DNA or RNA sequence can be represented as *r* = *b*_1_*b*_2_…*b*_*l*_, where *b*_*i*_ ∈ {*A, C, G, T, N* } or *b*_*i*_ ∈ {*A, C, G, U, N* }. Here, *A, G, C, T* and *U* represent the nitrogenous bases Adenine, Guanine, Cytosine, Thymine and Uracil, respectively. The letter *N* denotes an uncertain nucleotide, and *l* ∈ *ℕ* represents the total number of bases in *r*. We extract features from *r* by considering its substrings of length *k* (where 1 ≤ *k* ≤ *l*), also known as *k*-mers.

Each training instance consists of two reads in the edit-erring-READS or error-free-READS, examples of the training instances can be found in S1 Text. The features in reads are extracted using descriptors: **(i)** Fourier Transformation [48], **(ii)** Shannon’s and Tsallis’s entropy [48, 49], and **(iii)** Fickett’s score [50, 51]. Specifically, the features for a pair of reads with one or two base differences in a training instance may be identical. Therefore, features are only extracted from the absolute correct read (i.e., the first read in training instances) in order to avoid redundancy. These feature extraction methods are briefly depicted as follows:

i. Fourier Transformation. Sequences are converted into numeric representations by mapping each base to its atomic number (A:70, G: 78, C: 58, T or U: 66 and N: -1) [52]. Fourier Transformation converts the numerical sequence into a frequency domain representation using Discrete Fourier Transform (DFT). Here, Fast Fourier Transform (FFT) and a power spectrum analysis were conducted to extract features. This approach has been applied in several biological sequence analysis [53, 54], which can reveal hidden periodicity and repetitive elements in gene sequence. Mathematical exploration and examples of this approach can be found in [48].
ii. Shannon’s and Tsallis’s entropy. We first transformed each read sequence into its frequency representation by selecting a *k*-mer length of *k* = 1, 2, 3 and then computed both the absolute and relative frequencies of all possible *k*-mers. We then separately applied Shannon’s and Tsallis’s entropy to each *k*. These approaches capture the degree of disorder and uncertainty present in the sequence. Mathematical definitions and examples of these entropy approaches can be seen in [48].
iii. Fickett’s score involves the computation of four position-based values and four nucleotide-based content values from the sequence. The position indicator represents the degree of preference for each codon at one codon position over another, while the nucleotide-based content indicator measures the percentage of sequence per base. Then each indicator can be converted into a probability using the previous research output. The Fickett score is obtained by calculating the sum of probability multiplied by weight for each indicator. Detailed definitions and examples of this method refer to literature [51].

Additionally, we used the **(iv)** read counts and **(v)** characterised the error types and respective motifs as features. For instance, consider the two reads ACATG and ACGTG, the error is a substitution of C with G. Here, C-G is the error type, and CAT and CGT are the corresponding motifs. Similarly, for the two reads CGTG and ACGTG, the error is an insertion of A, the error type is represented as X-A, and the motifs are XA and AC. We define and normalise the feature vector *V* of error types or motifs as follows

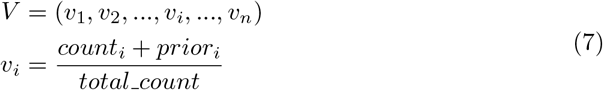

Here, *i* represents an error type, where *count*_*i*_ refers to the total number of occurrences of error type *i* and *total count* refers to the total number of all error types present in the data; *prior*_*i*_ is a small pre-defined value (e.g., 0.01) assigned to each item to avoid dividing by zero in cases where a certain error type or motif is not present in the data.

Before training, each type of feature is standardised separately by removing the mean and scaling to the unit variance using the pre-processing method “StandardScaler” of Scikit-learn [55]. To address class imbalance issue in the training data, we used the Synthetic Minority Over-sampling Technique (SMOTE) [56] with sampling performed by Imbalanced-learn [57].

### Model optimisation and prediction

The eXtreme Gradient Boosting (XGBoost) [12] is a well-established and efficient machine learning algorithm for classification. Optuna [58] is a framework that employs sampling and pruning heuristics to automatically discover optimal hyperparameter settings by conducting multiple trials. We chose XGBoost as our classifier and utilized Optuna to optimize the hyperparameters to achieve fast and accurate predictions.

We have pre-set some parameters for the classifier, including the tree method, regularisation term, number of estimators, and learning rate. A logistic regression was used to produce the probability for binary classification, and we aimed to maximize the test accuracy as the objective for selecting the best model via multiple trials (e.g., 20). For each task, noise2read utilized AutoML to create a new Optuna study object for training and selecting the best prediction model. For example, we trained and selected the four best models for predicting 1nt- and 2nt-based ambiguous, 1nt-based high ambiguous, and amplicon errors.

### Error correction for isolated nodes

After prediction, we restore all the edit-erring-READS to their normal state. We then adopted a third-party method Bcool [26] to deal with the errors contained in the isolated nodes, including many singletons, in the 1nt-edit-distance read graph. However, we keep only those corrected sequences by Bcool, which present in the original or in the first round of the corrected dataset without any genuine and ambiguous errors, to prevent generating non-existing new sequences.

### Evaluation criteria

#### Generation of the gold-standard wet-lab datasets with UMI-based ground truth

We have developed a novel approach for generating ground truth datasets, motivated by Mitchel’s method presented in literature [30]. Our method involves first correcting the Unique Molecular Identifier (UMI) errors using noise2read before clustering sequences with identical UMI tags. We did not attempt to correct those UMI sequences (if exist) which contain 2 base errors or more. The reason is that a UMI sequence is short with an extremely low probability containing two or more errors. Our detailed correction step for UMI sequences is just the construction of a 1nt-edit-distance read graph from the UMI sequences, and then the identification of those leaf nodes from the graph whose frequency is less than or equal to the maximum frequency threshold for the error reads and which also have a neighboring node with a higher frequency. Such a strategy might under-correct the errors in UMIs but will also avoid the over-correction issue for the UMIs. Additionally, within each UMI group, we retain one high-frequency read accounting for more than half of the group, which will avoid the impact when two or more sequences are bound to the same UMI tag.

Moreover, we observed that many raw reads contain a difference of more than three bases compared to their ground truth counterparts (see S16 Table) in [30]. For instance, in the dataset derived from SRR1543964, there are over 10,853 reads with a length of 238, each containing at least three base errors. Under the assumption in the introduction of this study, the likelihood of a read with a length of 238 having more than three base errors is relatively low, at only 0.00186, given a per-base error rate of 0.1%.

It is impossible to construct a dataset of absolute ground truth from a real sequencing dataset, even with the help of UMI tagging. Using UMI to create a ground truth from real sequencing data is equivalent to using UMI for error correction. To our knowledge, no substantiated evidence supports the rationality of altering three or more bases within a sequence to establish a ground truth. Through our above theoretical and edit distance analysis, we chose to drop those reads in each UMI cluster that have *>*2 errors to construct ground truth datasets. While this approach is not an ideal method in generating ground truth datasets, it can accurately capture a substantial portion of sequences in their actual states and those sequencing errors. We calculated and summarized the total numbers and the proportions of the sampled reads in the construction of UMI-based ground truth datasets used in this study and those in the benchmark study (see S20 Table). noise2read obtained a similar percentage of sampled reads by using the strict sampling strategy (only choosing one high frequency read in each UMI cluster and dropping all those reads in each UMI group that have *>*2 errors) in comparison with Mitchell’s method.

#### Generation of UMI-based simulated datasets from wet-lab datasets

We employed two different processes for generating UMI-based simulation datasets to evaluate the proposed method’s performance. The first process was designed to simulate single-end miRNA datasets similarly as worked in the literature [27]. Ideally, each unique read corresponds to a UMI. Therefore, we generate one unique UMI (numbers are used here instead of base sequences) for each unique sequence in the generated error-free read set. We then write these mimicked UMIs to each record in the edit-erring read set by mapping the sequence ID.

Taking motivations from the previous simulation approach [27], we introduced an innovative method to generate UMI-based simulation datasets that can be applied to a broader range of NGS datasets, extending beyond just miRNA sequencing data. The new approach consists of four steps: (1) Correcting a real sequencing dataset using a simplified version of noise2read which excludes complex feature extraction and machine learning components, opting instead for basic feature of frequency levels. This simplified approach is intended to prevent any potential bias in favour of noise2read when evaluating it with these simulated datasets. (2) Reads with counts below a predetermined threshold (e.g., 5) are filtered out to eliminate noise. This process generates a subset denoted as *S*0. Subsequently, those reads whose sequence counts surpass a predefined threshold (e.g., 30) within *S*0 are extracted, constituting an error-prone subset named *S*1, while the remaining reads form an error-free subset referred to as *S*2. (3) In this stage, reads are randomly sampled from *S*1 at predetermined error rates per read. These error-prone reads encompass instances with either 1 or 2 base errors, without overlaps. Subsequently, 1 or 2 base errors are randomly induced within these error-prone reads, resulting in simulated raw datasets termed *S*3. These settings in steps (2) and (3) will further avoid any potential bias favouring noise2read that might inadvertently arise during the first step of the simulation. (4) Combine *S*2 and *S*3 to create simulated datasets with errors, combine *S*1 and *S*2 to create simulated datasets of ground truth, and then create mimic UMIs for the simulated datasets using the same methodology above.

#### Metrics

To accurately evaluate the performance of error correction methods, we propose using confusion matrices at the read-level and base-level to measure changes in reads within the same UMI cluster rather than relying on the sequencing IDs generated by the instruments. The rationale is that for the constructed UMI-based datasets in this study, there is only one unique error-free sequence (of multiple occurrences) in each UMI cluster. Therefore, in a UMI cluster, we only need to compare the edit distance between every other unique read and this error-free read before and after error correction. Then, we are able to compute the confusion matrix using the relevant read count information before and after correction. More than half of the calculation time was saved this way. Otherwise, if we use the sequencing ID as the index, we must compare the edit distances twice for each group (same sequencing ID) of the raw, error-free, and corrected reads, even if they are the same. The absolute values of the true positives in each dataset are associated with the number of reads rather than the number of UMIs. The total number of positives of the actual condition equals the sum of the True positive and False negative. At the read level, a read is deemed erroneous if even a single base is incorrect. Conversely, a read is considered error-free only when all its bases are correct. TP defines the number of edit-erring reads perfectly modified after correction, and TN is the amounts of error-free reads without any changes after modification. While FP denotes the counts of error-free reads that are incorrectly adjusted by introducing new errors, FN represents the number of unchanged or wrongly fixed edit-erring reads. Similarly, at the base level, TP, TN, FP and FN concerned about the mistaken or accurate bases changing before and after correction.

Additionally, we employ the edit distance changes instead of the multiple sequence alignment (MLA) strategies used in [30] among raw, authentic and modified reads to get the confusion conditions as the MLA is highly time-consuming. Another reason is that MLA has more alternative alignment results since it compares three reads. In contrast, the edit-distance-based strategy only compares the ground truth read to its raw or corrected one, respectively. When counting the FN on the base level, we measure the absolute edit distance difference with the accurate read before and after correction.

Then, we derive the True Positive Ratio (TPR, a.k.a. recall or sensitivity), False Negative Ratio (FNR), True Negative Ratio (TNR), False Positive Ratio (FPR, a.k.a. fall-out), precision, gain and accuracy from the confusion matrix. TPR and FNR are the ratio of the number of edit-erring reads or bases correctly rectified and wrongly kept as negatives to the total number of actual edit-erring reads or bases, respectively. TNR and FPR are defined as the ratio of the number of error-free reads or bases correctly kept as negatives and wrongly rectified to the total number of actual error-free reads or bases, respectively. From the information theory perspective, TPR is the ratio of noise turning to signal; in contrast, FNR is the unconverted ratio of noise to signal. While FPR is the percentage of new noise introduced, TNR is the ratio of the original signal preserved.

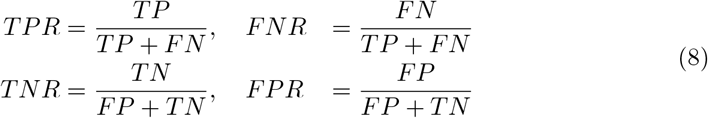

An ideal performance should achieve high TPR and TNR while keeping FNR and FPR low. Therefore, we can construct a cross-coordinate system where derived scores from the confusion matrix are assigned to each of the four directions. The four index values of TPR in the upper axis, FPR in the lower axis, TNR in the right axis and FNR in the left axis form a rectangle. The larger the overlapping area between the rectangle and the upper right quadrant, the better the performance. Therefore, we define a quantitative metric of the overlapping Area Difference (AD) to assess the performance as follows,

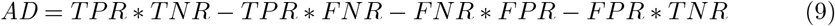

Moreover, we also calculate the Precision, Positive Gain and Accuracy denoted as follows to evaluate the correction performance at read-level or base-level. Precision evaluates the ratio of precise modifications among all the completed corrections and all the errors, while Positive Gain indicates the positive effect among all the real errors. The accuracy is the proportion of accurate modifications, including true positives and negatives, to the total number of reads or bases concerned.

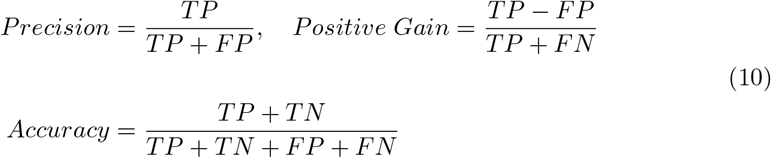

Furthermore, based on the read-level definition for the dataset of known ground truth, we classify reads into two categories: edit-erring and error-free. Then we measure the purity of the dataset using entropy defined as

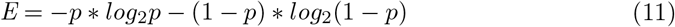

where *p* is the probability of randomly selecting one error-prone or error-free read from all sequences. The lower the dataset entropy, the fewer edit-erring reads exist in the dataset.

## Data availability

Generated datasets of UMI-based ground truth are available at D1-D8, their original sequencing data was obtained from SRR1543964-SRR1543971. Simulated ground truth datasets with mimic UMIs are available at D9-D13 with their original sequencing data downloaded from SRR12060401, SRR9077111 and SRR11092062. Simulated miRNA sequencing datasets with mimic UMIs are available at D14-D17. Processed SARS-Cov-2 data are available at D18-D19 with the corresponding source data available at SRR11092062. Monkeypox sequencing data D20-D21 were obtained from SRR22085311. Datasets of D22-D31 for the case study of isomiRs identification and SNPs profiling are available at ERR187525, ERR187844, ERR187542, ERR187668, ERR187759, ERR187669, ERR187622, ERR187665, ERR187892, ERR187500. Datasets for Adenine and cytosine base editor analysis are available at D32-D33 which are generated from Supplementary Table2 of the literature [44]. UMI-based ground-truth datasets (D34-D41 in this study) previously established in the literature [30] can be found at https://doi.org/10.6084/m9.figshare.11776413. S1 Table summarizes the information of datasets *D*1-*D*41 used in this study.

## Code availability

The algorithm, noise2read, developed in this study is packaged and released on the Python Package Index (PyPI) at https://pypi.org/project/noise2read/ and Bioconda at https://anaconda.org/bioconda/noise2read with source code publicly available at https://github.com/Jappy0/noise2read and documentation publicly available at https://noise2read.readthedocs.io/en/latest/.

## Supporting information

Supporting information, including Supplementary Figures 1-24 (presented in a single PDF), Supplementary Tables 1-20 (presented in a single PDF) and Supplementary Notes, are extended data supporting the results of this study.

**S1 Fig**. The example of two UMI clusters shows the number of low-frequency sequences with the corresponding smallest edit distance compared to the high-frequency sequences in the data set SRR1543971.

**S2 Fig**. The proportion of the UMI clusters with at least 60% of the erroneous reads caused by 1 or 2 base errors under the assumption that those low-frequency reads with an edit distance equal to or less than 4 may be erroneous reads caused by sequencing errors

**S3 Fig**. The example of five UMI clusters shows low-frequency sequences with the corresponding smallest edit distance compared to the high-frequency sequences in the data set SRR1543971.

**S4-S10 Figs**. Information gain visualisation for noise2read, Care, RACER, Bcool, Pollux, Lighter, Fiona and Coral on datasets D2-D10.

**S11 Fig**. The comparison results regarding Positive Rate (TPR), True Negative Rate (TNR), False Positive Rate (FPR), False Negative Rate (FNR), Area Difference (AD) at base-level between noise2read with seven other computational methods on the UMI-based wet-lab datasets D1-D8.

**S12 Fig**. The comparison results regarding Positive Rate (TPR), True Negative Rate (TNR), False Positive Rate (FPR), False Negative Rate (FNR), Area Difference (AD) at read-level between noise2read with seven other computational methods on the UMI-based wet-lab datasets D2-D8.

**S13-S17 Figs**. Comparison of True Positive Rate (TPR), True Negative Rate (TNR), False Positive Rate (FPR), False Negative Rate (FNR), Area Difference (AD), Precision, Gain and Accuracy at the base- and read-level, purity Entropy (E) and Non-frequent reads Entropy at the read-level between noise2read and Care, RACER, Bcool, Pollux, Lighter, Fiona and Coral on simulated datasets D9-D13.

**S18-S21 Figs**. Information gain visualisation for noise2read, Care, RACER, Bcool, Pollux, Lighter, Fiona and Coral on datasets D10-D13.

**S22-S24 Figs**. Scatter graphs compare the numbers of ambiguous isomiRs, exclusive isomiRs and known SNPs from miRNA sequencing datasets D23-D31 after error correction of the reads, while the Kepler plots provide additional visualisations of the data distribution.

**S1 Table**. The information of datasets D1-D33 used in this study.

**S2-S9 Tables**. The comparison results regarding TP, FP, FN, TN, Accuracy, Precision, Recall, Gain and Fall-out on the base- and read-level and the purity Entropy change (△*E*) and non-frequent reads’ information gain (△*H*) on the read-level between noise2read and other seven methods including Coral, RACER, Fiona, Lighter, Pollux, Bcool and Care on the UMI-based real sequencing datasets *D*1-*D*8.

**S10-S14 Tables**. The comparison results regarding TP, FP, FN, TN, Accuracy, Precision, Recall, Gain and Fall-out on the base- and read-level and the purity Entropy change (△*E*) and non-frequent reads’ information gain (△*H*) on the read-level between noise2read and other seven methods including Coral, RACER, Fiona, Lighter, Pollux, Bcool and Care on the simulated datasets *D*9-*D*13.

**S15 Table**. Evaluation comparison regarding TP, FP, FN, TN, Accuracy, Precision, Recall, Gain and Fall-out on the base- and read-level and the purity Entropy change (△*E*) and non-frequent reads’ information gain (△*H*) on the read-level between noise2read with and without enabling high ambiguous error prediction and miREC on miRNA simulation datasets *D*14-*D*17.

**S16 Table**. The number of reads pairs regarding edit distance between raw and ground truth datasets generated from SRR1543964 to SRR1543971 in literature [30].

**S17 Table**. The comparative performance between noise2read and the methods of Bless, Coral, Lighter, Reckoner, Sga, BFC, Pollux, Fiona, RACER, and Care on the UMI-based benchmark data sets *D*34-*D*37 established in the literature [30].

**S18 Table**. The comparative performance between noise2read and the methods of Bless, Coral, Lighter, Reckoner, Sga, BFC, Pollux, Fiona, RACER, and Care on the UMI-based benchmark data sets *D*38-*D*41 established in the literature [30].

**S19 Table**. The runtime in different stages and computational resources provided during error correction by noise2read on the data sets of *D*1-*D*41.

**S20 Table**. The total number of the sequences after sampling from the original data sets, as well as their percentage, when constructing UMI-based ground truth data sets in this study and the benchmark study, respectively.

**S1 Text. Supplementary Notes** for software commands used and examples of the training and predicting instances. (PDF)

## Acknowledgments

We would like to thank the computational resources provided by the UTS (University of Technology Sydney) eResearch High-Performance Computer Facilities.

## Author Contributions

**Conceptualization:** Jinyan Li, Pengyao Ping, Xuan Zhang.

**Data Curation:** Pengyao Ping, Xuan Zhang, Xinhui Cai, Hui Peng.

**Formal Analysis:** Pengyao Ping, Jinyan Li, Xuan Zhang, Tian Lan, Shuquan Su, Xinhui Cai, Hui Peng.

**Funding Acquisition:** Jinyan Li.

**Investigation:** Pengyao Ping, Xuan Zhang.

**Methodology:** Pengyao Ping, Jinyan Li, Xuan Zhang, Tian Lan, Shuquan Su.

**Project Administration:** Jinyan Li.

**Resources:** Pengyao Ping, Jinyan Li.

**Software:** Pengyao Ping.

**Supervision:** Jinyan Li, Wei Liu.

**Validation:** Pengyao Ping.

**Visualization:** Pengyao Ping, Shuquan Su, Xinhui Cai.

**Writing - Original Draft Preparation:** Pengyao Ping, Jinyan Li.

**Writing - Review & Editing:** Jinyan Li, Pengyao Ping, Wei Liu, Yi Pan.

## Declarations

The authors declare that they have no competing financial interests.

